# Plastid genomics of *Nicotiana* (Solanaceae): insights into molecular evolution, positive selection and the origin of the maternal genome of Aztec tobacco (*Nicotiana rustica*)

**DOI:** 10.1101/2020.01.13.905158

**Authors:** Furrukh Mehmood, Abdullah, Zartasha Ubaid, Iram Shahzadi, Ibrar Ahmed, Mohammad Tahir Waheed, Péter Poczai, Bushra Mirza

## Abstract

The genus *Nicotiana* of the family Solanaceae, commonly referred to as tobacco plants, are a group cultivated as garden ornamentals. Besides their use in the worldwide production of tobacco leaves, they are also used as evolutionary model systems due to their complex development history, which is tangled by polyploidy and hybridization. Here, we assembled the plastid genomes of five tobacco species, namely *N. knightiana, N. rustica, N. paniculata, N. obtusifolia and N. glauca*. *De novo* assembled tobacco plastid genomes showed typical quadripartite structure, consisting of a pair of inverted repeats (IR) regions (25,323–25,369 bp each) separated by a large single copy (LSC) region (86,510 –86,716 bp) and a small single copy (SSC) region (18,441–18,555 bp). Comparative analyses of *Nicotiana* plastid genomes showed similar GC content, gene content, codon usage, simple sequence repeats, oligonucleotide repeats, RNA editing sites and substitutions with currently available Solanaceae genomes sequences. We identified twenty highly polymorphic regions mostly belonging to intergenic spacer regions (IGS), which could be appropriate for the development of robust and cost-effective markers to infer the phylogeny of genus *Nicotiana* and family Solanaceae. Our comparative plastid genome analysis revealed that the maternal parent of the tetraploid *N. rustica* was the common ancestor of *N. paniculata* and *N. knightiana*, and the later species is more closely related to *N. rustica*. The relaxed molecular clock analyses estimated that the speciation event between *N. rustica* and *knightiana* appeared 0.56 Ma (HPD 0.65–0.46). The biogeographical analysis showed a south-to-north range expansion and diversification for *N. rustica* and related species, where *N. undulata* and *N. paniculata* evolved in North/Central Peru, while *N. rustica* developed in Southern Peru and separated from *N. knightiana*, which adapted to the Southern coastal climatic regimes. We further inspected selective pressure on protein-coding genes among tobacco species to determine if this adaptation process affected the evolution of plastid genes. These analyses indicated that four genes involved in different plastid functions, such as DNA replication (*rpo*A) and photosynthesis (*atpB*, *ndh*D and *ndhF*), came under positive selective pressure as a result of specific environmental conditions. Genetic mutations of the following genes might have contributed to the survival and better adaptation during the evolutionary history of tobacco species.

## 1. Introduction

The plant family Solanaceae consists of 98 genera and ∼ 2700 species (Olmstead et al., 2008; Olmstead & Bohs, 2007). This megadiverse family consists of herbaceous annual species to perennial trees with a natural distribution ranging from deserts to rainforests (Knapp et al., 2004). *Nicotiana* L. is the fifth largest genus in the family, comprising 76 species, which were subdivided into three subgenera and fourteen sections by Goodspeed (1954). The subgenera of *Nicotiana*, as proposed by Goodspeed (1954), were not monophyletic (Aoki & Ito, 2000; Chase et al., 2003), but most of Goodspeed’s sections were natural groups. The formal classification of the genus has been refined to reflect the growing body of evidence on *Nicotiana*, consisting of thirteen sections (Knapp, Chase & Clarkson, 2004). Most *Nicotiana* species are diploid (2n = 2x = 24) while allopolyploid species are also reported (Leitch et al., 2008). Phylogenetic studies have shown that these allopolyploids were formed 0.2 million (*N. rustica* L. and *N. tabacum* L.) to more than 10 million years ago (species of sect. *Suaveolentes*) (Clarkson et al., 2004; Leitch et al., 2008). Cultivated tobacco (*N. tabacum* L.), commonly grown for its leaves and an important economic and agricultural crop around the world (Occhialini et al., 2016), is a natural amphiploidy derived from two progenitors (Smith 1974). *Nicotiana* species, especially *N. tabacum*, are also used as model organisms in plant sciences and genetics (Zhang et al., 2011). The first complete chloroplast genome sequence was also published for this species (Shinozaki et al., 1986). Since the publication of this sequence, the structure and composition of chloroplast genomes has become widely utilized in identifying unique genetic changes and the evolutionary relationships of various groups of plants, while plastid genes have also been linked with important crop traits such as yield and resistance to various pest and pathogens (Jin & Daniell, 2015). Chloroplasts (cp) are large double membrane organelles with a genome size of 75-250 kb (Palmer, 1985). Proteins are used not only for photosynthesis but also for the synthesis of fatty acids and amino acids (Cooper, 2000). Angiosperm plastomes commonly contain ∼130 genes with up to 80 protein-coding, 30 transfer RNA (tRNA), and four ribosomal RNA (rRNA) genes (Daniell et al., 2016). The plastid genome exists in circular and linear forms (Oldenburg & Bendich, 2015) and the percentage of each form varies within plant cells (Oldenburg & Bendich, 2016). Circular formed plastomes have a typical quadripartite structure, with two inverted repeat regions (IRa and IRb), separated by one large single-copy (LSC) and one small single-copy (SSC) region (Palmer, 1985; Amiryousefi, Hyvönen & Poczai, 2018a; Abdullah et al., 2019b). Numerous mutational events occur in plastid genomes: variations in tandem repeats, insertion and deletions (indels), and point mutations, but inversions and translocations are also common (Jheng et al., 2012; Xu et al., 2015; Abdullah et al., 2019a). The plastid genome of angiosperms have a uniparental maternal inheritance (Daniell, 2007) but paternal inheritance has been recorded in a few gymnosperm species (Neale & Sederoff, 1989). The conserved organization of the plastid genome makes it extremely useful in exploring the phylogenetic relationships at various taxonomic levels (Ravi et al., 2008). Polymorphism in the chloroplast genome has been exploited to solve taxonomic issues, infer phylogeny and to investigate species adaptation to their natural habitats (Daniell et al., 2016). Genes in the plastid genome encode proteins and several types of RNA molecules, which play a vital role in functional plant metabolism, and can consequently undergo selective pressures. Most plastid protein-coding genes are under purifying selection to maintain their function, while positive selection might act on some genes in response to environmental changes. Complete plastid genome sequences are also useful tools in population genetics (Ahmad, 2014), species barcoding (Nguyen et al., 2017) transplastomic (Waheed et al., 2011, 2015) and conservation of endangered species (Wambugu et al., 2015).

Here, we assembled the plastid genome of five *Nicotiana* species and compared their sequences to gain insight into the chloroplast genome structure of the genus *Nicotiana*. We also inferred the phylogenetic relationship of genus *Nicotiana* and investigated the selection pressures acting on protein-coding genes, then identified mutational hotspots in the *Nicotiana* plastid that might be used for the development of robust and cost-effective markers in crop breeding or taxonomy.

## 2. Materials and Methods

### 2.1. Chloroplast genomes assembly and annotation

Illumina sequence data of *Nicotiana knightiana* L. (13.1 Gb, accession number SRR8169719*), N. rustica (*15.5 Gb, SRR8173839*), N. paniculata* (35.1 Gb, SRR8173256*), N. obtusifolia (*23 Gb, SRR3592445) and *N. glauca (*12.5 Gb, SRR6320052) were downloaded from the Sequence Read Archive (SRA). The chloroplast genome sequence contigs were selected by performing the BWA alignment with default settings (Li & Durbin, 2009) using *Nicotiana tabacum* (GenBank accession number: NC_001879) as a reference. Geneious R8.1 *de novo* assembler (Kearse et al., 2012) was used to order the selected contigs for final assembly. The genome sequence was annotated using GeSeq (Tillich et al., 2017) and CPGAVAS2 (Shi et al., 2019). Following *de novo* annotation, start/stop codons and the position of introns were manually inspected and curated. The tRNA genes were verified by tRNAscan-SE version 2.0 with default settings (Lowe & Chan, 2016) and Aragorn version 1.2.38 (Laslett & Canback, 2004). Circular genome maps were drawn with OGDRAW v1.3.1 (Greiner, Lehwark & Bock, 2019). The average coverage depth of *Nicotiana* species plastid genomes was determined by mapping all reads to *de novo* assembled plastid genomes with BWA (Li & Durbin, 2009) visualized with Tablet (Milne et al., 2009). Novel *Nicotiana* plastid genomes were deposited in NCBI and the assigned accession numbers are shown in Table 1.

### 2.2. Comparative genome analysis and RNA editing prediction

Novel plastid genome sequences were compared through multiple alignments using MAFFT v7 (Katoh & Standley, 2013). Every part of the genome, such as intergenic spacer regions (IGS), introns, protein-coding genes, and ribosomal RNAs and tRNAs, was considered for comparison. Each part was extracted and used to determine nucleotide diversity in DnaSP v6 (Rozas et al., 2017). Substitution, transition and transversion rates were also calculated compared to the *N. tabacum* reference using Geneious R8.1 (Kearse et al., 2012). Structural units of the plastid genome (LSC, SSC and IR) were individually aligned to determine the rate of substitutions and to further search for indels using DnaSP v6. The expansion and contraction of inverted repeats and their border positions were compared for ten selected *Nicotiana* species using IRscope (Amiryousefi, Hyvönen and Poczai, 2018b). The online software PREP-cp (Putative RNA Editing Predictor of Chloroplast) was used with default settings to determine putative RNA editing sites (Mower, 2009) and the codon usage and amino acids frequencies were determined by Geneious R8.1 software (Kearse et al., 2012).

### 2.3. Repeats analyses

Microsatellites repeats within the plastid genomes of five *Nicotiana* species were detected using MISA (Beier et al., 2017) with the minimal repeat number of 7 for mononucleotide repeats, 4 for di- and 3 for tri-, tetra-, penta- and hexanucleotide SSRs. We also used REPuter software (Kurtz, 2002) with the following parameters: minimal repeats size was set to 30 bp, Hamming distance to 3, minimum similarity percentage of two repeats copies up to 90%, maximum computed repeats numbers to 500 bp for scanning and visualizing forward (F), reverse (R), palindromic (P) and complementary (C) repeats. Tandem repeats were found with the tandem repeats finder using default parameters (Benson, 1999).

### 2.4. Synonymous (*K*_s_) and non-synonymous (*K*_a_) substitution rate analysis

The synonymous (*K*_s_) and non-synonymous (*K*_a_) substitution were analyzed using the chloroplast genome of *Nicotiana tabacum* as reference for all *de novo* assembled *Nicotiana* plastid genomes. For this purpose, protein-coding genes were extracted from *Nicotiana* plastomes, then aligned with the corresponding genes of *S. dulcamara* as a reference using MAFFT (Katoh & Standley, 2013) and analyzed using DnaSP software (Rozas et al., 2017). We further assessed the impact of positive selection using additional codon models to estimate the rates of synonymous and nonsynonymous substitution. The signs of positive selection were further assessed using fast unconstrained Bayesian approximation (FUBAR) (Murrell et al., 2013) and the mixed effects model of evolution (MEME) (Murrell et al., 2012) as implemented in the DATAMONKEY web server (Delport et al., 2010). Sites with cut-off values of PP < 0.9 in FUBAR were considered as candidates to have evolved under positive selection. Out of all analyses performed in DATAMONKEY, the most suited model of evolution for each data set, directly estimated on this web server, was used. In addition, the mixed effects model of evolution (MEME), a branch-site method incorporated in the DATAMONKEY server, was used to test for both pervasive and episodic diversifying selection. MEME applies models variable ω across lineages at individual sites, restricting ω to be ≤ 1 in a proportion p of branches and unrestricted at a proportion (1 − p) of branches per site. Positive selection was inferred with this method for a P value < 0.05.

### 2.5. Phylogenomic analyses

Plastid genome sequences from the genus *Nicotiana* were selected from Organelle Genome Resources of NCBI, accessed on 21.2.2019. We included all available plastid genome sequences of tobacco species in our analysis and added *de novo* assembled sequences while *S. dulcamara* was used as an outgroup. For the species included in our analysis, coding alignments were constructed from the excised plastid genes using MACSE (Ranwez et al., 2011), including the following seventy-five protein coding genes: *atp*A, B, E, F, H, I; *ccs*A; *cem*A; *clp*P; *mat*K; *ndh*A, B, C, D, E, F, G, H, I, J, K; *pet*A, B, D, G, J, L, N; *psa*A, C, I, J; *psb*A, B, C, D, E, F, I, L, M, N, T, Z; *rbc*L; *rpl*2, 7, 14, 16, 19, 20, 22, 23, 32, 33, 36; *rpo*A, B, C1; *rps*2, 3, 4, 7, 8, 11, 14, 15, 16, 18, 19; *ycf*2, 3, 4. For phylogenetic analysis we used a matrix of protein-coding genes of twelve species with a concatenated matrix length of 75,449 bp. The best fitting model (GY+F+I+G4) was determined by ModelFinder (Kalyaanamoorthy et al., 2017) as implemented in IQ-TREE according to the Akaike information criterion (AIC), and Bayesian information criterion (BIC). Maximum likelihood (ML) analyses were performed with IQ-TREE (Nguyen et al., 2015) using the ultrafast bootstrap approximation (UFBoot; Hoang et al., 2018) with 1,000 replicates and the SH-like approximate likelihood ratio test (SH-aLRT), also with 1,000 bootstrap replicates, and TreeDyn was used for further enhancement of phylogenetic tree analysis (Dereeper et al., 2008; Lemoine et al., 2019).

Relative divergence times were estimated for the species *N. rustica* and putative parental species using BEAST v.1.8.4 (Drummond et al., 2012), applying GTR + I + G rate substitution to the protein-coding plastid gene matrix. A Yule speciation tree prior and a relaxed uncorrelated clock-model that allows rates to vary independently along branches (Drummond et al., 2006) were used, with all other parameters set to default. The median time split between the *S. dulcamara* and *N. undulata* (mean = 25 Myr; standard deviation = 0.5) was used as a temporal constraint to calibrate the BEAST analyses derived from the Solanaceae-wide phylogeny of Särkinen et al. (2013) and the Time Tree of Life (Kumar et al. 2017). Uncertainty regarding these dates was incorporated by assigning normal prior distributions to the two calibration points (Couvreur et al., 2008; Evans et al., 2014). Four independent BEAST runs were conducted, each with 10 million generations, sampling every 10,000 generations. Convergence of all parameters was assessed in Tracer 1.5 (Rambaut et al., 2014) and 10% of each chain was removed as burn-in. The Markov chains were combined in LogCombiner 1.7.2. (Drummond et al., 2012) to calculate the maximum clade credibility tree.

We defined six biogeographical areas based on Köppen-Geiger climatic and further biogeographic evidence and distributions: (A) Colombian/Ecuadorian mountain range mixed equatorial (Af), monsoon (Am) and temperate oceanic climate (Cfb), (B) Northern Peruvian mountain range with tropical savanna climate (Aw), (*C*) Central Peru with equatorial climate (*Af*), (*D*) Coastal Peru with cold semi-arid and desert climate (*Bsk*, *BWk*), (*E*) Peruvian Mountain range with humid subtropical/oceanic highland climate (*Cwb*), (*F*) Bolivian/Chilean alpine/mountain range with mixed semi-arid cold (*Bsk*, *BWk*) and humid subtropical climate (*Cwa*). These areas were used in the Bayesian Binary Method (BBM) model implemented in RASP (Yu, Blair & He, 2019) to investigate the biogeographic history of the selected four *Nicotiana* species. BBM infers ancestral area using a full hierarchical Bayesian approach and hypothesizes a special “null distribution”, meaning that an ancestral range contains none of the unit areas (Ronquist 2004). The analysis was performed on the BEAST maximum clade credibility tree using default settings, i.e. fixed JC + G (Jukes-Cantor + Gamma) with null root distribution. Ancestral area reconstruction for each node was manually plotted on the BEAST tree using pie charts. Species distributions were determined from data stored in the Solanaceae Source Database (http://solanaceaesource.org/) and Global Biodiversity Information Facility (GBIF) (https://www.gbif.org/).

## 3. RESULTS

### 3.1. Characteristics of *Nicotiana* plastid genomes

Five *Nicotiana* species chloroplast genomes were assembled and the lengths of these plastid genomes were: *Nicotiana knightiana* (155,968 bp)*, Nicotiana rustica* (155,849 bp)*, Nicotiana paniculata* (155,689 bp)*, Nicotiana obtusifolia* (156,022 bp) and *Nicotiana glauca* (155,917 bp). Further details of the characteristics of the assembled plastid genomes are summarized in Table S1. The coverage of assembled plastid genomes was 811× *Nicotiana knightiana*, 1,951× *Nicotiana rustica*, 1,032× *Nicotiana paniculata*, 1,412× *Nicotiana obtusifolia* and 327× *Nicotiana glauca*. The GC content of IR regions were highest (43.2%) followed by LSC (35.9%) and SSC (32.1%) (Table S1). The high GC content of IR was due to high GC content of the tRNAs (52.9%) and rRNAs (55.4%) genes.

*De novo* assembled *Nicotiana* plastid genomes had 134 unique genes, whereas eighteen genes were duplicated in the IR region (Table S2, Fig.1). Out of 134 genes, 86 were protein-coding genes, 37 were tRNA genes and 8 were rRNA genes. Among 18 duplicated genes in IR region, 7 were protein-coding, 7 were tRNA genes and 4 were rRNA genes. 18 intron-containing genes were present in the plastome of *Nicotiana* species. The *rps12* gene is a trans-spliced gene, its 1^st^ exon existing in the LSC region while the 2^nd^ and 3^rd^ exons are in the IR region.

**Figure 1.**
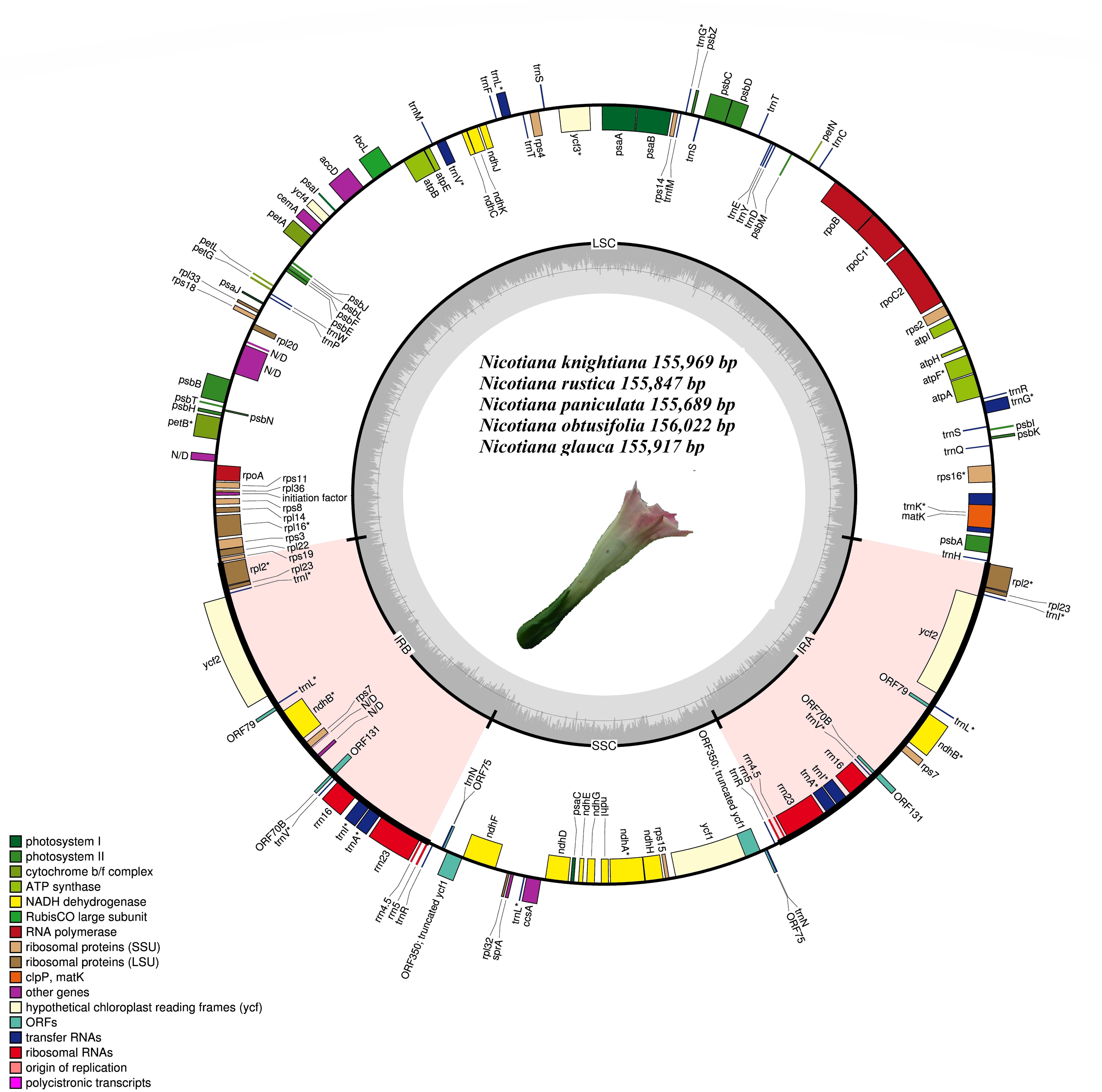
Chloroplast genome map of *Nicotiana knightiana, Nicotiana rustica, Nicotiana paniculata, Nicotiana obtusifolia* and *Nicotiana glauca.* Genes that lie outside the circle are transcribed clockwise while the genes that transcribed counterclockwise are inside the circle. Different colors indicate the genes belonging to various functional groups. GC and AT content of genome are plotted light grey and dark, respectively, in the inner circle. Large single copy (LSC), inverted repeat A (IRa), inverted repeat B (IRb) and small single copy (SSC) are shown in the circular diagram. Inverted repeat regions are highlighted with *cinderella* color.

### 3.2. Comparative analyses, codon usage and RNA editing sites

The nucleotide composition of *Nicotiana* species was compared, and all genomes had similar nucleotide composition indicating high synteny in the LSC, SSC, IR and CDSs but also in non-coding regions. Detailed comparison of the base composition is shown in Table S3. A high percentage of hydrophobic amino acids were encoded in *Nicotiana* plastid genomes, while acidic amino acids were present at lower rates. The amino acids are AT rich sequences as compared to GC (Fig. 2A). Relative synonyms codon usage (RSCU) and frequency of amino acid revealed that leucine is the most abundant and cysteine was the least encoded amino acid in these genomes (Fig S1). The codon usage revealed a high frequency of codons with A/T at 3^rd^ codon position as compared to C/G at 3^rd^ codon position (Table S4).

**Figure 2.**
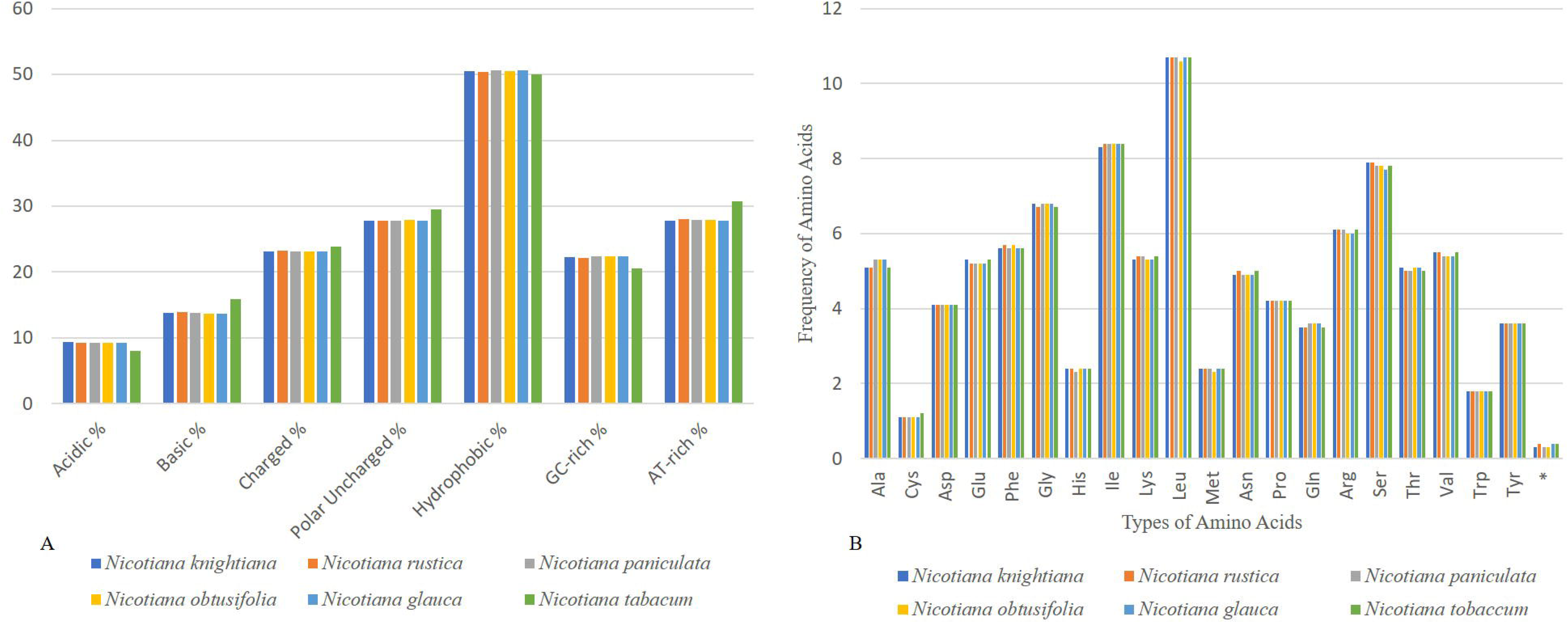
(A) Comparison of amino acid groups in *Nicotiana knightiana, Nicotiana rustica, Nicotiana paniculata, Nicotiana obtusifolia, Nicotiana glauca.* (B) Comparison of amino acid frequency in *Nicotiana knightiana, Nicotiana rustica, Nicotiana paniculata, Nicotiana obtusifolia, Nicotiana glauca*.

The number of predicted RNA editing sites using PREP-cp varied between 34 and 37, distributed among fifteen genes (see Table S3). Among these genes, *ndhB* (9) possessed the most of these sites, followed by *ndhD* (6-8) and *rpoB* (4). The *ndhD* gene revealed a fraction of variation among species: *N. knightiana, N. rustica* and *N. paniculata* having six RNA editing sites whereas seven were observed in *N. obtusifolia* and eight in *N. glauca*. Most of the RNA editing sites were C to U edits on the first and second base of the codons, but the frequency of second base codon edits was much higher. The conversions from serine to leucine were the most frequent and these changes helped in the formation of hydrophobic amino acids, i.e. valine, leucine and phenylalanine (Table S5).

### 3.3. IR contraction and expansion

The LSC/IR and IR/SSC border positions of *Nicotiana* plastid genomes were compared (Fig 3) using IRscope. The length of the IR regions was similar, ranging from 25,331bp to 25,436bp showing some expansion. The endpoint of the Solanaceae JLA (IRa/SSC) is characteristically located upstream of the *rps*19 and downstream of the *trn*H-GUG, which was confirmed in *Nicotiana*. In *N. tomentosiformis*, the IR expanded to partially include *rps*19, creating a truncated ψ*rps*19 copy at JLA, which was thought to be missing from the entire *Nicotiana* clade (Amiryousefi, Hyvönen & Poczai, 2018a). In this species the IR region has expanded to include 60 bp of *rps*19. The extent of the IR expansion to *rps*19 varied from 2 to 60 bp and the end point seems to be conserved to the following intergenic spacer region. Furthermore, *inf*A, *ycf*15, and a copy of *ycf*1 located on the JSB were detected as pseudogenes. The position of *ycf1* in the IRb/SSC region varied. It left a 36 bp pseudogene in *N. knightiana*, *N. rustica* and *N. glauca*, 33 bp pseudogene in *N. obtusifolia* and a 72 bp one in *N. paniculata*.

**Figure 3.**
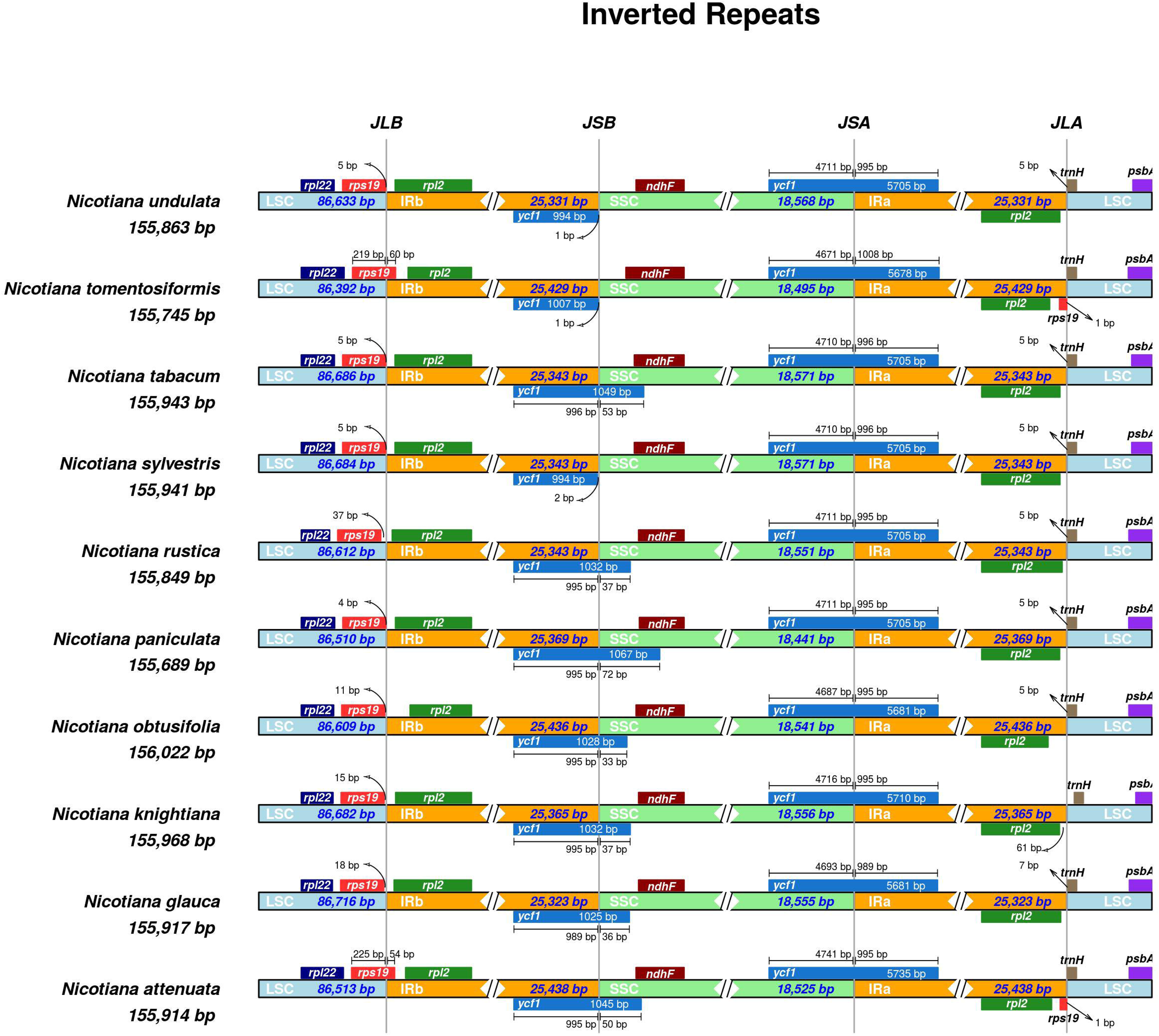
Comparison of the border positions of LSC, SSC and IR among the five *Nicotiana* chloroplast genomes. Positive strand transcribed genes are indicated under the line while the genes that are transcribed by negative strands are indicated above the line. Gene names are expressed in boxes, and the lengths of relative regions are showed above the boxes. The number of bp (base pairs) that are written with genes reveal the part of the genes that exists in the region of chloroplast or away from region of chloroplast i.e. bp written with *ycf1* indicate that sequences exist in that region of the plastid genome.

### 3.4. Non-synonymous (K_a_) and synonymous (K_s_) substitution rate analysis

Synonymous/non-synonymous substitutions ratio is widely used as an indicator of adaptive evolution or positive selection (Kimura, 1979). We have calculated the K_s_, K_a_ and K_a_/K_s_ ratio for 77 protein-coding genes for five selected *Nicotiana* species using *S. dulcamara* as a reference (Table S6). Among the analyzed genes, 31 had K_s_=0, 19 had K_a_=0, and 39 genes had both K_s_ and K_a_=0 values. Of the investigated genes, 21 showed a K_a_/K_s_ ratio of more than 0.5. Eight of these genes (*atpF, psaA, ycf4, psbB, infA, ndhB, rpl32* and *ccsA)* had a K_a_/K_s_ ratio greater than 0.5 for one species, *ycf1* had K_a_/K_s_ greater than 0.5 for two species, while *atpA, rps2, rpoB, rps12, ycf2, ndhG* had K_a_/K_s_ greater than 0.5 for three species whereas genes *rpoC1, atpB, rpoA, ndhD* had K_a_/K_s_ ratio more than 4 species and *rpoC2* and *ndhF* had K_a_/K_s_ ratio for all species. We selected the genes *atp*B, *rpo*A, *ndh*D, *ndh*F, *rpo*C1 and C2 for further analysis using FUBAR and MEME. FUBAR estimates the number of nonsynonymous and synonymous substitutions at each codon given a phylogeny, and provides the posterior probability of every codon belonging to a set of classes of ω (including ω = 1, ω < 1 or ω > 1) (Murrell et al., 2013). MEME estimates the probability for a codon to have undergone episodes of positive evolution, allowing the ω ratio distribution to vary across codons and branches in the phylogeny. This last attribute allows identification of the proportion of codons that may have been evolving neutrally or under purifying selection, while the remaining codons can also evolve under positive selection (Murrell et al., 2012). The two models indicated positive selection on the codons only found in *atp*B, *rpo*A, *ndh*F and *rpo*A (Table 1). Thus, the methods described suggested six amino acid replacements altogether as candidates for positive selection, of which three were fixed in all *Nicotiana*, and three were restricted to diverse groups of species (see Table 1).

### 3.5. Repetitive sequences in novel *Nicotiana* plastid genomes

Repeat analysis performed with MISA revealed high similarity in chloroplast microsatellites (cpSSRs) ranging from 368 to 384 among tobacco species. The majority of the SSRs in these plastid genomes were mononucleotide rather than trinucleotide or dinucleotide. The most dominant of the SSRs were A/T motifs mononucleotides, and in dinucleotides AT/TA motifs were the second most predominant. Mononucleotide SSRs varied from 7-17 units repeats; dinucleotide SSRs from 4-5-unit repeats while other SSRs types were present mainly in 3-unit repeats. Mostly the SSRs existed in LSC, in comparison to IR and SSC (Fig 4) (Table S7). REPuter software was used to identify and locate forward (F), reverse (R), palindromic (P), and complementary (C) repeats in all the species of *Nicotiana.* In the plastomes of five *Nicotiana* species, we found 117 oligonucleotide repeats: 25 in *N. knightiana*, 23 in *N. rustica*, 21 in *N. paniculata*, 23 in *N. obtusifolia*, 25 in *N. glauca*. Forward (F) and palindromic repeats were present in large numbers as compared to others in all species: 11 (44%) (F) and 14 (56%) (P) in *N. knightiana*, 14 (60%) (F) and 9 (39%) (P) in *N. rustica*, and 12 (57%) (F) and 9 (42%) (P) in *N. paniculata*, 14 (56%) (F) and 11 (44%) (P) in *N. obtusifolia*, 9 (39%) (F) and 11 (52%) (P) in *N. glauca*. The size of oligonucleotide repeats varied from 30-65 bp, and many of these repeats were 30-35 bp in length. The LSC region held most of the identified oligonucleotide repeats as compared to SSC and IR. The LSC region contained 13 in *N. knightiana*, 11 in *N. rustica*, 14 in *N. paniculata*, 15 in *N. obtusifolia* and 17 in *N. glauca*. In plastid genome regions, the repeats existed mostly in IGS, followed by CDS and intronic regions (Fig. 5) (Table S8). The number of tandem repeats varies from 24-27 between these *Nicotiana* species. The IGS region contains the most tandem repeats followed by the CDS region. The size of these repeats varied between 20 to 88 among *Nicotiana* (Fig. 6).

**Figure 4.**
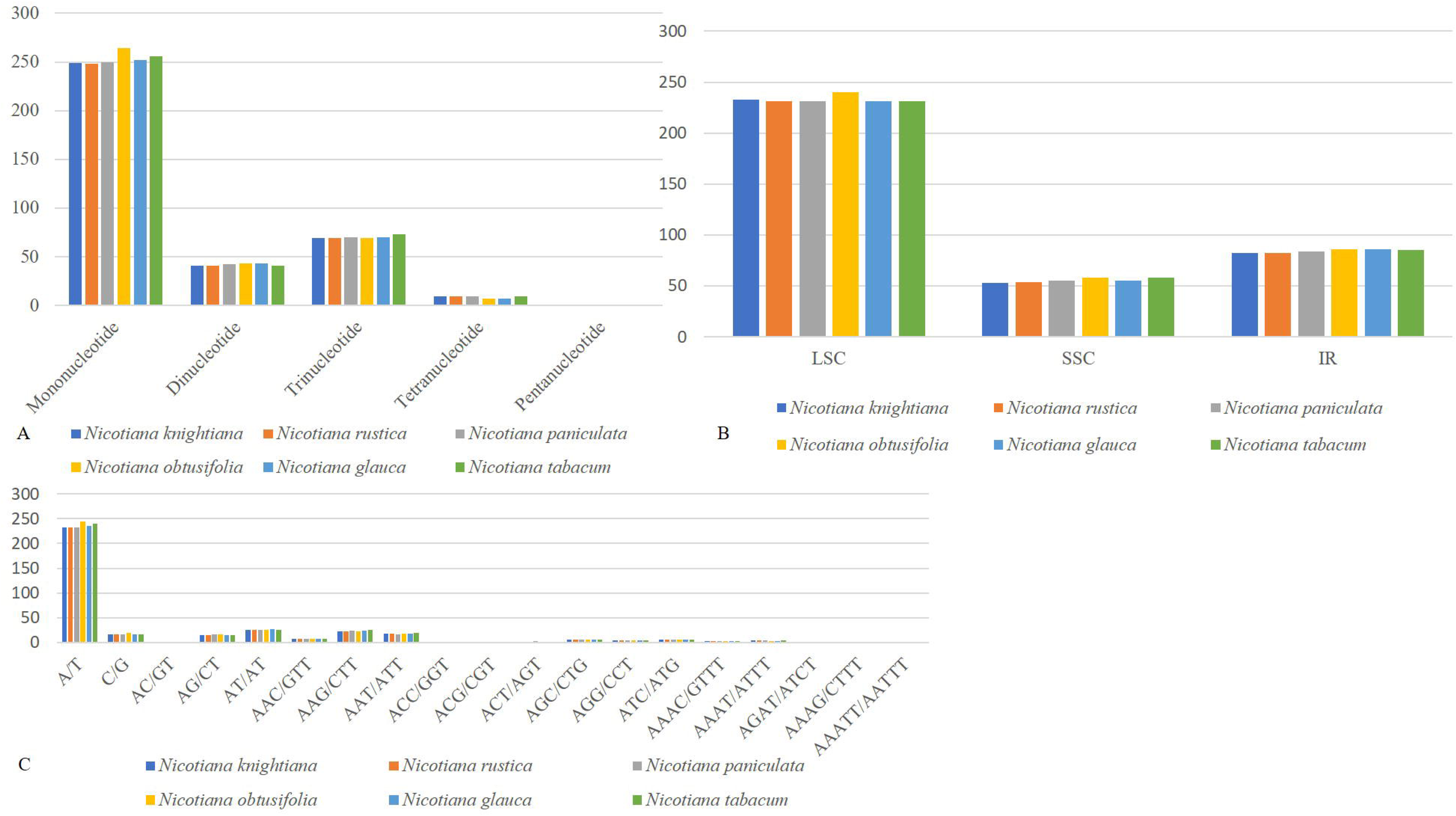
Comparison of microsatellite repeats among *Nicotiana knightiana, Nicotiana rustica, Nicotiana paniculata, Nicotiana obtusifolia, Nicotiana glauca. (A)* Indicate numbers of various types of microsatellites present in the plastid genome of *Nicotiana* species. *(B)* Distribution of SSRs in different regions of the plastid genome of *Nicotiana* species. *(C)* SSRs motifs distribution in different regions of the plastid genome of *Nicotiana* species.

**Figure 5.**
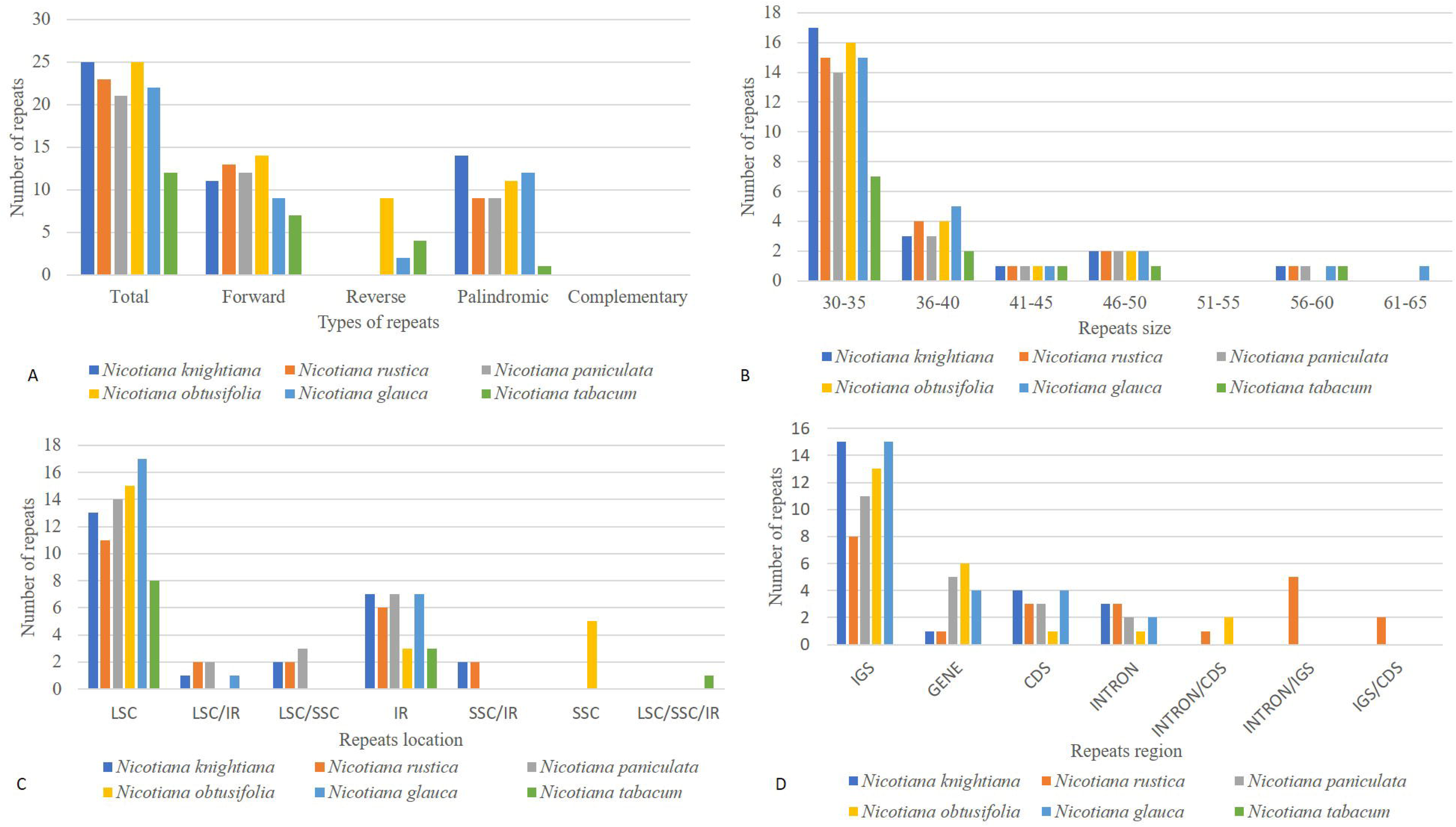
(A). Indication of various kinds of oligonucleotide repeats exist in all *Nicotiana species* (B). Indicate repeats that exist range of size i.e. 30–35 indicate numbers of repeats within the size vary from 30 and 35. (C). Indicate number of repeats exist in separate areas of plastid genome. LSC: Large single copy, SSC: small single copy, IR: inverted repeat region, LSC/SSC: one copy of LSC and another in SSC, LSC/IR: one copy of LSC and another in SSC, IR/SSC: one copy of IR and another in SSC, LSC/SSC/IR: one copy of LSC, one in SSC and another in IR. (D). Indicate number of repeats in different regions of plastid genome. IGS: Intergenic spacer region, CDS: coding DNA sequences, Intron: intronic regions, IGS/Intron: one copy of intergenic spacer region and another in intronic regions. Intron/CDS: one copy intron region and another in CDS regions. IGS/CDS: intergenic spacer region copy of repeat and one more in coding regions.

**Figure 6.**
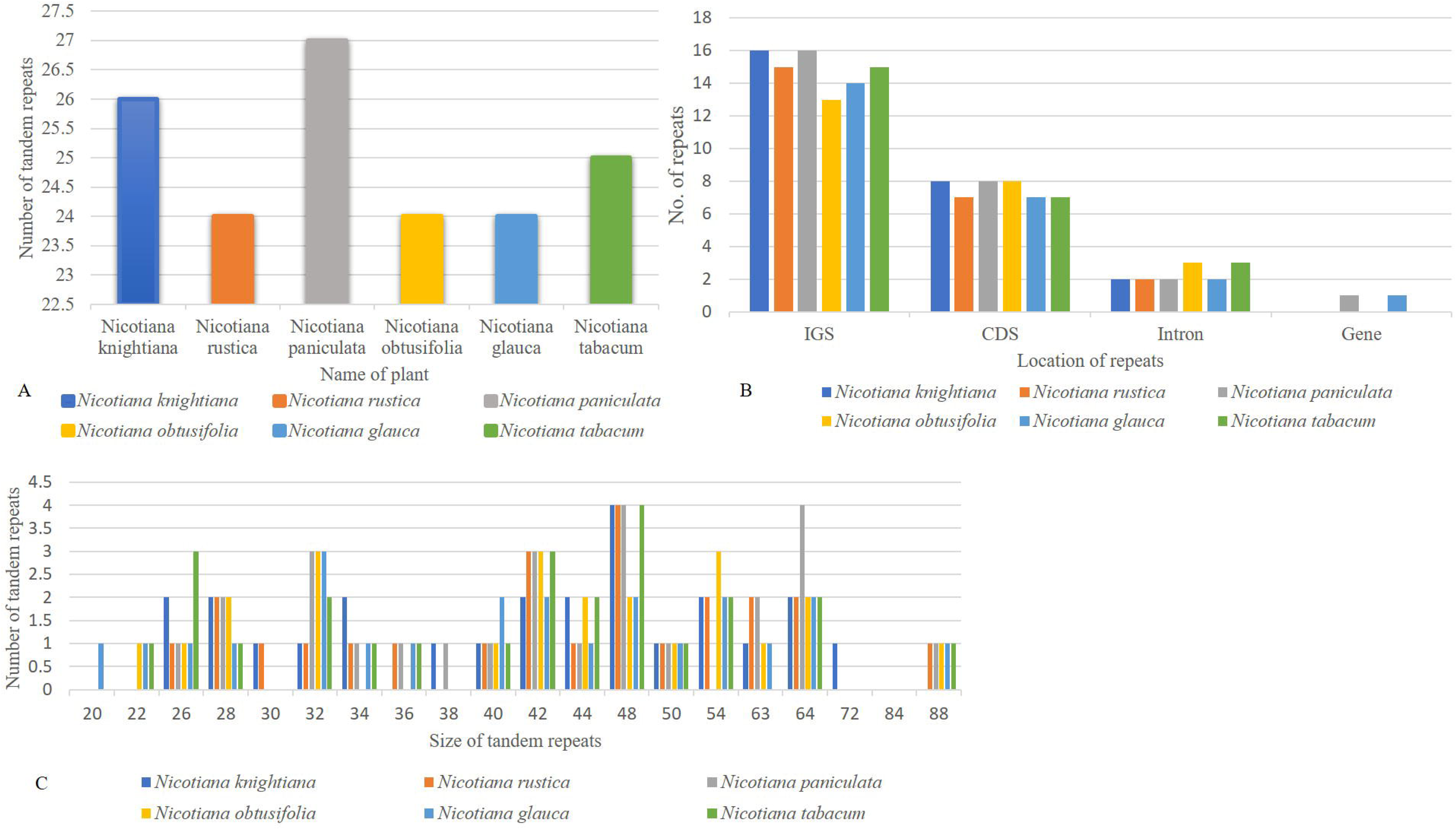
Comparison of tandem repeats among *Nicotiana knightiana, Nicotiana rustica, Nicotiana paniculata, Nicotiana obtusifolia, Nicotiana glauca. (A)* Number of tandem repeats in the chloroplast genome of *Nicotiana knightiana, Nicotiana rustica, Nicotiana paniculata, Nicotiana obtusifolia, Nicotiana glauca. (B)* Location and number of tandem repeats in the plastid genome of *Nicotiana knightiana, Nicotiana rustica, Nicotiana paniculata, Nicotiana obtusifolia, Nicotiana glauca. (C)* Number, size, distribution of tandem repeats across the plastid genome of *Nicotiana knightiana, Nicotiana rustica, Nicotiana paniculata, Nicotiana obtusifolia, Nicotiana glauca*

### 3.6. Single nucleotide polymorphism and insertion/deletion analyses in *Nicotiana*

We investigated substitution types in the five plastomes of *Nicotiana* species (one IR removed), using *Nicotiana tabacum* as a reference. *Nicotiana knightiana* (786), *Nicotiana rustica* (775), *Nicotiana paniculata* (861)*, Nicotiana obtusifolia* (847) and *Nicotiana glauca* (509) substitutions were seen in the whole plastid genome. The types of substitutions exhibited among *Nicotiana* species were similar. Most of the conversions were A/G and C/T in comparison to other SNPs (single nucleotide polymorphism) (Table 2). Ts/Tv ratio were as follows: *Nicotiana knightiana* LSC (1.5), SSC (0.968) and IR (1.047), *Nicotiana rustica* LSC (1.496), SSC (0.978) and IR (1), *Nicotiana paniculata* LSC (1.461), SSC (0.886) and IR (0.833), *Nicotiana obtusifolia* LSC (1.097), SSC (1.020) and IR (1.194), *Nicotiana glauca* LSC (0.924), SSC (0.819) and IR (0.783) (Table S9). The substitutions in different regions of these genomes are *Nicotiana knightiana* contains 560 (LSC), 43 (IR) and 183 (SSC) SNPs, *Nicotiana rustica* contains 599 (LSC), 32 (IR) and 183 (SSC) substitutions, *Nicotiana paniculata* has 630 (LSC), 33 (IR) and 198 (SSC) substitutions, *Nicotiana obtusifolia* consists of 671 (LSC), 68 (IR) and 210 (SSC) substitutions while *Nicotiana glauca* has 327 (LSC), 82 (IR) and 100 (SSC). Insertions and Deletions (indels) were also examined using DnaSP in all regions of the chloroplast genome. In total, *Nicotiana knightiana* (110), *Nicotiana rustica* (107), *Nicotiana paniculata* (116)*, Nicotiana obtusifolia* (143) and *Nicotiana glauca* (113) indels were found. The LSC region held the majority of the indels, followed by SSC, whereas IR contained minimum indels (Table 3).

### 3.7. Divergence hotspot regions in *Nicotiana*

The CDS, intron and IGS regions of the whole plastid genome of five *Nicotiana* species were compared to discover polymorphic regions (mutational hotspots). High polymorphism was found in intronic regions (average π=0.167) in comparison to IGS (π=0.031) and CDS regions (average π=0.002). Among *Nicotiana* species, the nucleotide diversity values varied from 0 (*ycf3*) to 0.306 (*rps12 intron* region) (Fig. 7). Here, 20 highly polymorphic regions were determined that might be used as potential makers to reconstruct the phylogeny for identifying *Nicotiana* species (Table 4).

**Figure 7.**
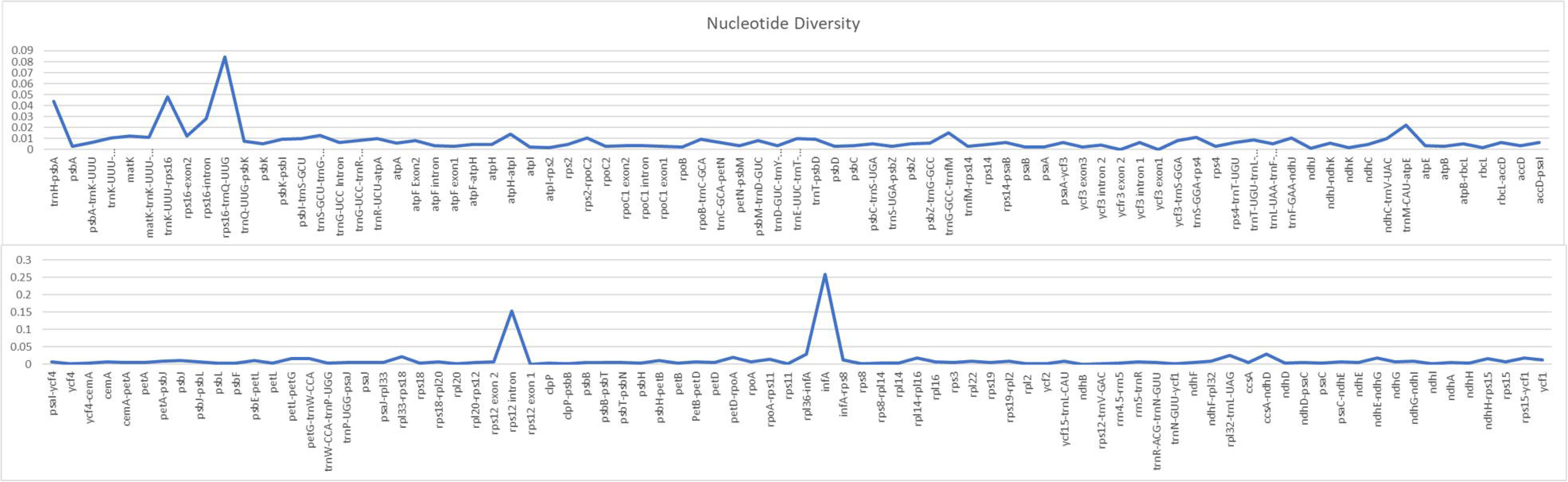
Nucleotide diversity of various regions of the chloroplast genome among *Nicotiana* species. The X-axis indicate the chloroplast regions and Y-axis indicate the nucleotide diversity.

### 3.8. Phylogenomic analyses

Phylogenetic analysis within *Nicotiana* plastid genomes were reconstructed with the maximum likelihood method, based on selected and concatenated protein-coding genes. Our phylogenetic analyses resulted in a highly resolved tree (Fig 8), with almost all clades recovered having maximum branch support values. After the elimination of indels, the tree was reconstructed based on alignment size of 75,449 bp with the best fitting model GY+F+I+G4 (Fig 8). We further concentrated on the species phylogeny of *N. rustica* and putative parental species where relative divergence times were estimated using a relaxed uncorrelated clock implemented in BEAST. This analysis found that the divergence of *N. undulata* appeared 5.36 (highest posterior density, HPD 6.38–4.43) million years ago (Ma), while *N. paniculata* diverged 1.17 Ma (HPD 2.18–0.63) followed by the most recent split of *N. rustica* and *N. knightiana* 0.56 Ma (HPD 0.65–0.46). This analysis showed that the *Nicotiana* species included in the analysis are not older than the end of the Pliocene and that most subsequent evolution must have occurred in the Pleistocene. The timing of these lineage splits, in addition to the current distributions of four closely related species, were used to infer the progression of migratory steps in RASP (Fig 9). The most recent common ancestor (MRCA) area illustrated a dispersal event for *N. paniculata* in Northern (B) and Southern Peru (E) and the vicariance of *N. knightiana* in Coastal Peru (D). The overall dispersal pattern of the examined species showed a south-to-north expansion pattern from Central Peru to Colombia and Ecuador (*N. rustica*) to Bolivia (*N. undulata*).

**Figure 8.**
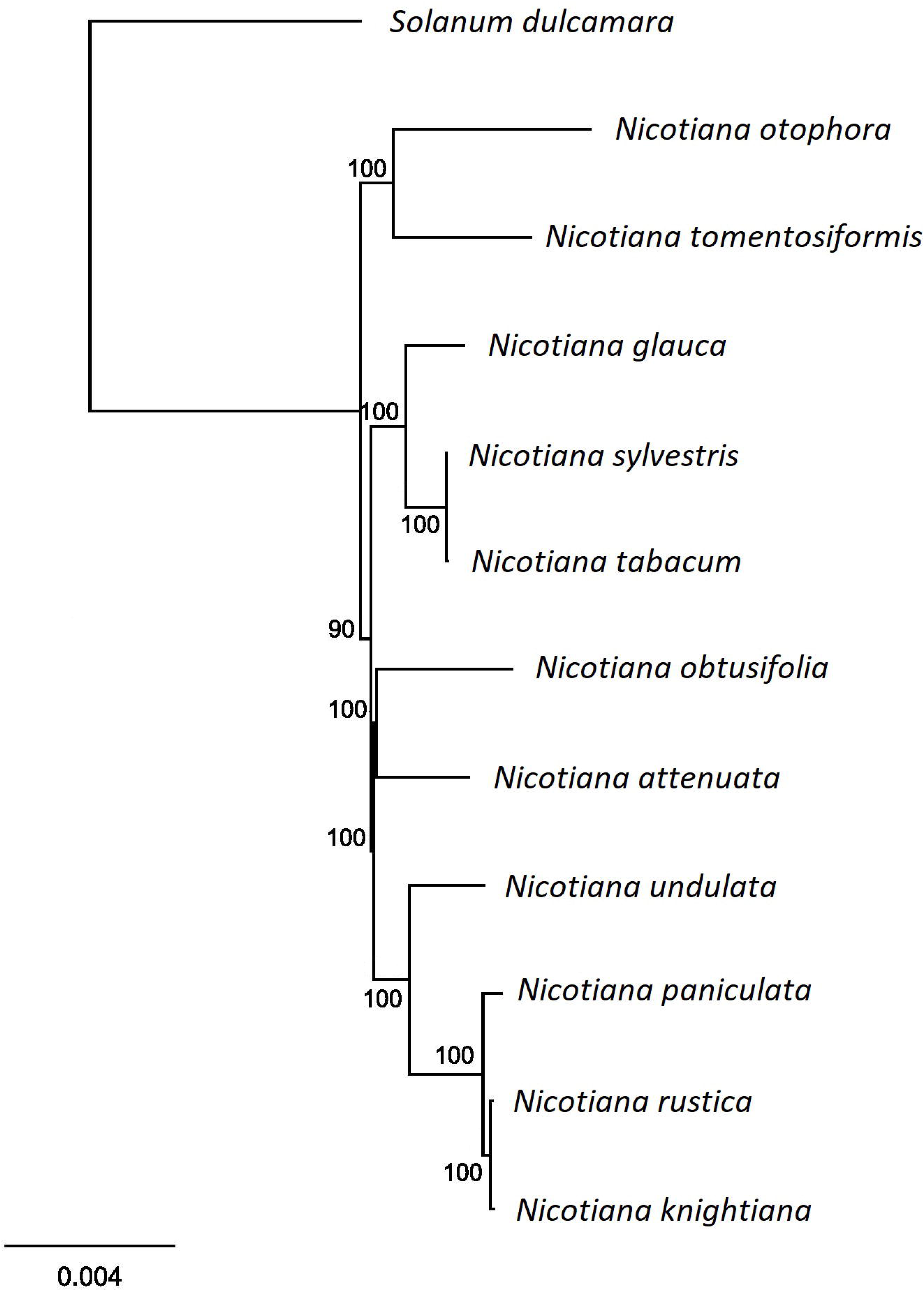
Maximum likelihood (ML) tree was reconstructed based on seventy-five protein coding plastid genes of eleven *Nicotiana* species and *Solanum dulcamara* as an outgroup. Bootstrap support values are shown above or below the nodes.

**Figure 9.**
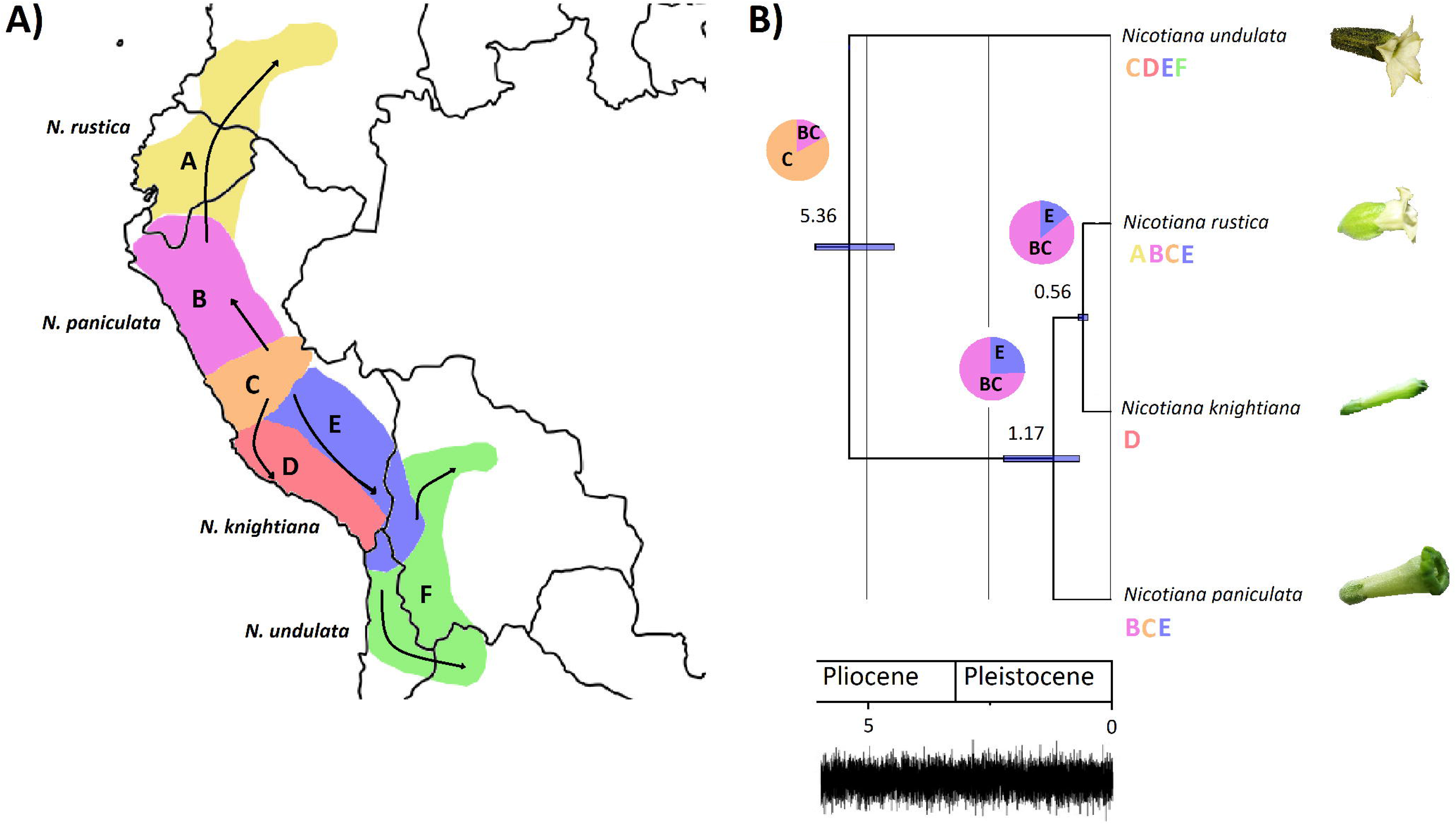
Plastome phylogeny and biogeography of the tetraploid *Nicotiana rustica* and related species. A) Map showing the six biogeographic areas used to infer the biogeographic history of the *Nicotiana rustica* in South America. Arrows illustrate the dispersal events inferred from the biogeographic analysis. Geographical distribution for each terminal is indicated using the biogeographic regions subdivision. The most probable ancestral area is figured at each node of the phylogenetic tree. Pie-charts represent relative probabilities of ancestral states at each node. B) Node-calibrated Bayesian maximum clade credibility tree with 95% highest posterior density (HPD) interval for node ages presented as horizontal bars and mean values are displayed above each node. All nodes have PP 0.97 and BS 87%. Trace plot of the combined chains showing the sampled joint probability and the convergence of the chains.

## 4. Discussion

### 4.1. Molecular evolution of *Nicotiana* plastid genomes

We compared five chloroplast genomes of *Nicotiana* species, which revealed similar genomic features. These comparative analyses produced an insight into the phylogeny and evolution of *Nicotiana* species. The GC content of the *Nicotiana* species referred to above were similar to those of other *Nicotiana* species (Sugiyama et al., 2005; Yukawa, Tsudzuki & Sugiura, 2006) i.e. the GC content in the IR is high, which might be a result of the existence of ribosomal RNA (Qian et al., 2013; Cheng et al., 2017; Zhao et al., 2018). The genome organization, gene order and content and of these *Nicotiana* species were also similar for *N. slyvestris* and *N. tabacum* (Sugiyama et al., 2005; Yukawa, Tsudzuki & Sugiura, 2006). The intron plays an important role in the regulation of gene expression (Xu et al., 2003). The *trn*K intron is important because it expresses an unusual form of a group II intron derived from a mobile group of mitochondrial-like intron open reading frames (ORFs) (Hausner et al., 2006). As in the plastid genomes of many land plants, the abundance of A/T content at 3^rd^ base of codons was reported due to high concentration of A/T nucleotides in the whole plastid genome (Menezes et al., 2018).

The plastomes of land plants have conserved structure but diversity prevails at the border position of LSC/SSC/IR of the genome. The size range of LSC, SSC and IR varies between the plastid genomes of species that advances to alterations in several genes and leads to the deletion of one copy of a gene or duplication of functional and non-functional genes of different sizes (Menezes et al., 2018; Saina et al., 2018). In the current study, all these ten *Nicotiana* species showed similarities with some variation as compared to the *Nicotiana tabacum:* in all these plants except *Nicotiana tomentosiformis* which have 60 bp in IRb region, the *rps19* gene is present entirely in the LSC region but in *Nicotiana tabacum rps19* gene extended 5bp in the IR region. The fluctuations at the border positions of various regions of the plastid genome might be helpful in determining the evolution of species (Menezes et al., 2018). Liu et al., (2018) reported that the similarities at the junction regions may be useful in explaining the relationship between the species and that those plants which have a high level of relatedness show minimal fluctuations at the junctions of the chloroplast genome. The resemblance at junctions reveals a close relationship between the *Nicotiana* species.

Repeats in the chloroplast genome are useful in evolutionary studies and play a vital role in genome arrangement (Zhang et al., 2016). Here, we detected that the mononucleotide repeats (A/T), and trinucleotide SSRs (ATT/TAA) were present in large amounts in all the species of *Nicotiana*, which may be a result of the A/T rich proportion of chloroplast genome. A similar result was also reported in *Nicotiana otophora* (Asaf et al., 2016). In all the species of *Nicotiana*, the LSC region contained a greater amount of SSRs in comparison to SSC and IR, which has also been demonstrated in other studies of angiosperm plastomes (Shahzadi et al., 2019; Mehmood et al., 2019). When less genomic resources are available for revealing divergence hotspot regions, the oligonucleotide repeats might be utilized as a substitute for identifying polymorphic regions (Ahmed et al., 2012; Ahmad, 2014). The current study results of oligonucleotide repeats are similar to the previously reported results of *Nicotiana* species and other angiosperm plastome studies (Asaf et al., 2016; Yang et al., 2019). Thus, the presence of both the high divergence regions in IGS and oligonucleotide repeats suggest that these regions are suitable for the development of markers to demonstrate phylogenetics relationships.

To understand the molecular evolution, it is important to know about the nucleotide substitution rates (Muse & Gaut, 1994). LSC and SSC regions are more prone to substitutions and indels whereas the IR regions are more conserved in the chloroplast genome (Ahmed et al., 2012; Abdullah et al., 2019b). Our results also showed similar results in that the IR region is mostly conserved, and most of the substitutions occurs in the LSC and SSC regions. Thus, the ratio (Ts/Tv) was equal to or more than 1. Similar results were shown in the chloroplast genome of *Dioscorea polystachya* (Yam) (Cao et al., 2018).

Divergence hotspot regions of the plastid genome could be used to develop accurate, robust and cost-effective molecular markers for population genetics, species barcoding and evolutionary based studies. (Ahmed et al., 2013; Ahmad, 2014; Nguyen et al., 2017). Previously, in several studies, polymorphic loci were identified based on comparisons of chloroplast genome to provide information about suitable loci for the development of molecular markers (Choi, Chung & Park, 2016; Li et al., 2018; Menezes et al., 2018). We found 20 polymorphic regions such as *infA, rps12 intron, rps16-trnQ-UUG* which have 0.25942, 0.15275, 0.08451 nucleotide diversity respectively that were more polymorphic than frequently used markers such as *rbcL*, and *matK*. These regions could be suitable markers for population genetics and phylogenetic analyses, especially in the genus *Nicotiana*.

### 4.2. Positive selection on *Nicotiana* plastid genes

Plants have evolved complex physiological and biochemical adaptations to adjust and adapt to a variety of environmental stresses. *Nicotiana*, originating in South America, has spread to many regions of the world and members of the genus have successfully adapted to harsh environmental conditions to survive. This great variation in their distributional range induced distinctive habits and morphology in the inflorescence and flowers, indicative of the physiological specialization to the area where they evolved. Desert ephemeral *Nicotiana* species are short while subtropical perennials have tall and robust habits with variable inflorescences ranging from pleiochasial cymes to solitary flowers and diffuse panculate-cymose mixtures. For example, members of *Nicotiana* section *Suaveolentes* Goodsp. evolving in isolation faced several cycles of harsh climate change. In Australia, the native range of the species, a predominantly warm and wet environment went through intensive aridification (Poczai, Hyvönen & Symon, 2011). Throughout this climate change and increasing central aridification, many species either retreated to the wetter coastline or adapted to and still survive in this hostile inland environment (Bally et al., 2018). Tobacco plants also developed specialized biosynthetic pathways and metabolites, such as nicotine, which serve complex functions for ecological adaptations to biotic and abiotic stresses, most importantly serving as a defense mechanism against herbivores (Xu et al. 2017). Therefore, *Nicotiana* is a rich reservoir of genetic resources for evolutionary biological research, since several members of the genus went through changing climatic events and adopted to environmental fluctuations.

The patterns of synonymous (K_s_) and non-synonymous (K_a_) substitution of nucleotides are essential markers in evolutionary genetics defining slow and fast evolving genes (Kimura, 1979). K_a_/K_s_ values >1, =1, and <1 indicate positive selection, natural evolution and purifying selection, respectively (Lawrie et al., 2013), while a minimal ratio of K_a_/K_s_ (<0.5) in many genes represents purifying selection working on them. Many proteins and RNA molecules encoded by the chloroplast genomes are under purifying selection since they are involved in important functions of plant metabolism, self-replication and photosynthesis and therefore play a pivotal role in plant survival (Piot et al., 2018). Departure from the main purifying selection in case of plastid genes might happen in response to certain environmental changes when advantageous genetic mutations might contribute to survival and better adaptation. The K_a_/K_s_ ratios in our analysis for *Nicotiana* species indicated changes in selective pressures. The genes *atp*B, *ndh*D, *ndh*F, *rpo*A, *rps*2 and *rps*12 had greater K_a_/K_s_ value (> 1), possibly due to positive selective pressure as a result of specific environmental conditions. This has been conclusively supported by an integrative analysis using Fast Unconstrained Bayesian AppRoximation (FUBAR) and Mixed Effects Model of Evolution (MEME) methods, which identified the set of positively selected codons in case of *atp*B, *ndh*D, *ndh*F and *rpo*A (Table 5), but provided no further evidence for *rps*2 and *rps*12.

These genes are involved in different plastid functions, such as DNA replication (*rpo*A) and photosynthesis (*atpB*, *ndh*D and *ndhF*). The *rpo*A gene encodes the alpha subunit of PEP, which is believed to predominantly transcribe photosynthesis genes (Hajdukiewicz, Allison & Maliga, 1997). The transcripts of plastid genes encoding the PEP core subunits are transiently accumulated during leaf development (Kusumi et al., 2011), thus the entire *rpo*A polycistron is essential for chloroplast gene expression and plant development (Zhang et al., 2018). The housekeeping gene *atp*B encodes the β-subunit of the ATP synthase complex, which has a highly conserved structure that couples proton translocation across membranes with the synthesis of ATP (Gatenby, Rothstein & Nomura, 1989), which is the main source of energy for the functioning of plant cells. In chloroplasts, linear electron transport mediated by PSII and PSI produces both ATP and NADPH, whereas PSI cyclic electron transport preferentially contributes to ATP synthesis without the accumulation of NADPH (Peng & Shikanai, 2011). Chloroplast NDH monomers are sensitive to high light stress, suggesting that the *ndh* genes encoding the NAD(P)H dehydrogenase (NDH) may also be involved in stress acclimation through the optimization of photosynthesis (Casano, Martín & Sabater, 2001; Martin et al., 2002; Rumeau, Peltier & Cournac, 2007). During acclimation to growth light environments, many plants change biochemical composition and morphology (Terashima et al., 2005). The highly responsive regulatory system controlled by cyclic electron transport around PSI could optimize photosynthesis and plant growth under naturally fluctuating light (Yamori, 2016). When the demand for ATP is higher than that for NADPH (e.g., during photosynthetic induction, at high or low temperature, at low CO_2_ concentration, or under drought), cyclic electron transport around PSI is likely to be activated (Yamori, 2016; Yamori & Shikanai, 2016). Thus, positive selection acting on ATP synthase and NAD(P)H dehydrogenase encoding genes is probably evidence for adaptation to novel ecological conditions in *Nicotiana*.

These findings might be also supported by our observation that RNA editing sites occurred frequently in *Nicotiana ndh* genes (Table S3). It has been shown that *ndh*B mutants under lower air humidity conditions or following exposure to ABA present a reduction in the photosynthetic level, likely mediated through stomatal closure triggered under these conditions (Horvath et al., 2000). Therefore, a protein structure modification resulting from a loss or decrease in RNA editing events could affect adaptations to stress conditions or cause other unknown changes (Rodrigues et al., 2017). Previous studies have demonstrated that abiotic stress influences the editing process and consequently plastid physiology (Nakajima & Mulligan, 2001). Alterations in editing site patterns resulting from abiotic stress could be associated with susceptibility to photo-oxidative damage (Rodrigues et al., 2017) and indicate that *Nicotiana* species experienced abiotic stresses during their evolution, which resulted in positive selection of some of the plastid genes. Up to this point, positive selection has rarely been detected in chloroplast genes except for *clp*P1 (Erixon & Oxelman, 2008), *ndh*F (Peng et al., 2011), *mat*K (Hao, Chen & Xiao, 2010) and *rbc*L (Kapralov et al., 2011). However, a recent study by Piot et al. (2018) showed that one-third of the plastid genes in 113 species of grasses (Poaceae) evolved under positive selection. This might indicate that positive selection might be overlooked among diverse groups of plant taxa.

### 4.3. Phylogenetic relationships and the origin of tetraploid *Nicotiana rustica*

Our comparative plastid genome analysis revealed that the maternal parent of the tetraploid *N. rustica* was the common ancestor of *N. paniculata* and *N. knightiana*, and the later species is more closely related to *N. rustica*. The relaxed molecular clock analyses estimated that the speciation event between *N. rustica* and *N. knightiana* appeared ∼0.56 Ma (HPD 0.65–0.46) in line with previous findings (Sierro et al., 2018). Comparative analysis of the genomes of four related *Nicotiana* species revealed that *N. rustica* inherited about 41% of its nuclear genome from its paternal progenitor, *N. undulata*, the rest from its maternal progenitor, the common ancestor of *N. paniculata* and *N. knightiana* (Sierro et al., 2018), which has also been confirmed by our study. It has been shown that *N. rustica* and in fact all *Nicotiana* tetraploids, except species included in section *Suaveolentes*, originated from a doubling of the diploid chromosome for the genus. Thus, they should be regarded as natural allopolyploids (Leitch et al., 2008). We also revealed that *N. knightiana* is more closely related to *N. rustica* than *N. paniculata*, which can be further corroborated by the distribution of indels highlighted in the present study. The biogeographical analysis carried out suggests that *N. undulata* and *N. paniculata* evolved in North/Central Peru, while *N. rustica* developed in Southern Peru and separated from *N. knightiana*, which adapted to the Southern coastal climatic regimes. Positively selected plastid genes with functions such as DNA replication (*rpo*A) and photosynthesis (*atpB*, *ndh*D and *ndhF*) might have been associated with successful adaptation to, for example, a coastal environment.

However, our results should be regarded as tentative, as our survey excludes several broad ecological variables from testing, including variation in salinity, island versus mainland, and East versus West of the Andes. We aim to highlight that many potential environmental variables might be highly correlated with speciation processes, as has been demonstrated in the same region for another Solanaceae group in the tomato clade (*Solanum* sect. *Lycopersicon*), where amino acid differences in genes associated with seasonal climate variation and intensity of photosynthetically active radiation were correlated with speciation processes (Pease et al., 2016). Another example of rapid adaptive radiation from the family is the genus *Nolana* L.f., where several clades gained competitive advantages in water-dependent environments by succeeding and diverging in Peru and Northern Chile (Dillon et al., 2009). In the case of *N. rustica* and related species we assume that diversification was driven by the ecologically variable environments of the Andes. Our molecular clock analysis provides recent species diversification in the Pleistocene and Pliocene while substantial climatic transitions in Peru predate these events. For example, the uplift of the central region of the Andes and the formation of the Peruvian coastal desert ended (∼14 – 150 Mya; Hoorn et al., 2010; Gerreaud et al., 2010) before the geographical and ecological expansion of *N. rustica* and related parental species.

The dispersal of *N. rustica* and related species shows a south-to-north range expansion and diversification which has been suggested by phylogenies of other plant and animal groups in the Central Andes (Picard, Sempere & Plantard, 2008; Lueber and Weigend, 2014). Based on the south-to-north progression scenario, habitats located at high altitudes were first available for colonization in the south, recently continuing to northward. Erosion and orogenic progression caused dispersal barriers of species colonizing these high habitats to diversify in a south-to-north pattern, frequently following allopatric speciation. Thus, for taxonomic groups currently residing throughout a large portion of the high Andes, a south-to-north speciation pattern is expected (Doan, 2003). In this case the most basal species (*N. undulata*) has more southern geographic ranges, and the most derived species (*N. rustica*) has more northern geographic ranges except for *N. knightiana*, which presumably colonized the coastal range of Peru. Although the four *Nicotiana* species examined show overlaps in their distribution, it is probable that speciation was caused by fragmentation of populations during the glacial period (see Simpson, 1975). Utilizing fewer chloroplast loci for phylogenetic analyses of plant species may limit the solution of phylogenetic relationships, specifically at low taxonomic levels (Hilu & Alice, 2001; Majure et al., 2012). Previously, genus *Nicotiana* was subdivided into 13 sections using multiple chloroplast markers, i.e. *trnL* intron and *trnL-F* spacer*, trnS-G* spacer and two genes*, ndhF and matK* (Clarkson et al., 2004). Recently, inference of phylogeny based on complete chloroplast genomes has provided deep insight into the phylogeny of certain families and genera (Henriquez et al., 2014; Amiryousefi, Hyvönen & Poczai, 2018a; Abdullah et al., 2019a). Here, we reconstructed a phylogenetic tree among eleven species of genus *Nicotiana* that belong to nine sections (Clarkson et al., 2004) based on 75 protein-coding genes by using *S. dulcamara* as an outgroup which attests the previous classification of genus *Nicotiana* with high bootstrapping values. Species of each section are well resolved whereas the *N. tabacum* of section *Nicotiana* and *N. sylvestris* of section Sylvestres show close resemblance. The *N. paniculata* and *N. knightiana* belong to section Paniculatae but here did not appear on the same node. This revealed that further data is required to elucidate the phylogenetic relationship among these two species. Overall, our phylogenetic analyses support the previous classification of genus *Nicotiana*, but enrichment of chloroplast genomic resources can provide further insight into the phylogeny of the genus *Nicotiana*.

## 5. Conclusion

In the present study, we assembled, annotated and analyzed the whole cp genome sequence of five *Nicotiana* species. The genomic structure and organization of their chloroplast genome was like those of previously reported Solanaceae plastomes. Divergences of LSC, SSC and IR region sequences were identified, as well as the distribution and location of repeat sequences. The identified mutational hotspots sequences could be utilized as potential molecular markers to investigate phylogenetic relationships in the genus. As we demonstrated in our study to elucidate the maternal genome origins of *N. rustica*, our results could provide further help in understanding the evolutionary history of tobaccos.

## Acknowledgements

We thank Kenneth Quek for editing the manuscript.

## Authors contributions

Furrukh Mehmood: Conceptualization, Genome assembly and annotation, Data analysis, Data interpretation, prepared figures and tables, Manuscript drafting and editing.

Abdullah: Genome annotation, Data analysis, Data interpretation, Manuscript drafting.

Zartasha Ubaid: Data analysis, Data interpretation, Manuscript drafting.

Iram Shehzadi: Data analysis, Data interpretation, Manuscript drafting.

Ibrar Ahmed: Conceptualization, Manuscript editing.

Mohammad Tahir Waheed: Conceptualization, Manuscript editing.

Péter Poczai: Supervision, carried out selection tests and phylogenetic analysis, prepared figures and tables, authored and reviewed drafts of the paper, approved the final draft.

Bushra Mirza: Supervision, authored or reviewed drafts of the paper, approved the final draft.

---

**Table 1.** List of amino acid replacements and results of positive selection tests on codons underlying these replacements.

**Table 2.** Comparison of substitution in *Nicotiana* species

**Table 3.** Distribution of indels in *Nicotiana* chloroplast genome

**Table 4.** Mutational hotspots among *Nicotiana* species

