## Supplementary material for "Plastid genomics of *Nicotiana* (Solanaceae): insights into molecular evolution, positive selection and the origin of the maternal genome of Aztec tobacco (*Nicotiana rustica*)": Table 1

**Table 1.** Summary statistics of de novo assembled Nicotiana plastid genomes

| **Characteristics** | | ***Nicotiana knightiana*** | ***Nicotiana rustica*** | ***Nicotiana paniculata*** | ***Nicotiana obtusifolia*** | ***Nicotiana glauca*** |
| --- | --- | --- | --- | --- | --- | --- |
| GenBank Accession Nr. | | BK010737 | BK010738 | BK010741 | BK010739 | BK010740 |
| Size (base pair; bp) | | 155,968 | 155,849 | 155,689 | 156,022 | 155,917 |
| LSC length (bp) | | 86,682 | 86,612 | 86,510 | 86,609 | 86,716 |
| SSC length (bp) | | 18,551 | 18,551 | 18,441 | 18,541 | 18,555 |
| IR length (bp) | | 25,364 | 25,343 | 25,369 | 25,436 | 25,323 |
| Number of genes | | 134 | 134 | 134 | 134 | 134 |
| Protein-coding genes | | 86 | 86 | 86 | 86 | 86 |
| tRNA genes | | 37 | 37 | 37 | 37 | 37 |
| rRNA genes | | 8 | 8 | 8 | 8 | 8 |
| Duplicate genes | | 18 | 18 | 18 | 18 | 18 |
| GC content | Total (%) | 37.1% | 37.8% | 37.9% | 37.8% | 37.8% |
|  | LSC (%) | 35.9% | 35.9% | 35.9% | 35.9% | 36% |
|  | SSC (%) | 32.1% | 32.1% | 32.2% | 31.9% | 32.1% |
|  | IR (%) | 43.2% | 43.2% | 43.2% | 43.2% | 43.2% |
|  | CDS (%) | 38.2% | 38.2% | 38.2% | 38.1% | 38.2% |
|  | rRNA (%) | 55.3% | 55.4% | 55.3% | 55.2% | 55.1% |
|  | tRNA (%) | 52.9% | 52.9% | 52.9% | 52.9% | 52.9% |
|  | All gene % | 39.8% | 39.9% | 39.8% | 39.7% | 39.8% |
| Protein coding part (CDS) (%bp) | | 52.6% | 50.2% | 51.7% | 51.6% | 38.2% |
| All gene (%bp) | | 70.43% | 70.72% | 72.04% | 71.99% | 71.96% |
| Non-coding region (%bp) | | 29.57% | 29.28% | 27.95% | 28.01% | 28.04% |
