## Supplementary material for "Plastid genomics of *Nicotiana* (Solanaceae): insights into molecular evolution, positive selection and the origin of the maternal genome of Aztec tobacco (*Nicotiana rustica*)": Table 2

**Table 2.** Comparison of substitution in *Nicotiana* species

| **Types** | ***Nicotiana knightiana*** | ***Nicotiana rustica*** | ***Nicotiana paniculata*** | ***Nicotiana obtusifolia*** | ***Nicotiana glauca*** |
| --- | --- | --- | --- | --- | --- |
| A/G | 222 | 219 | 245 | 244 | 110 |
| C/T | 226 | 223 | 237 | 250 | 128 |
| A/C | 105 | 104 | 117 | 153 | 97 |
| C/G | 40 | 39 | 50 | 52 | 34 |
| G/T | 130 | 129 | 135 | 148 | 87 |
| A/T | 63 | 61 | 77 | 102 | 53 |
| Total | 786 | 775 | 861 | 847 | 509 |
| **Location wise distribution** | | | | | |
| **LSC** | 560 | 559 | 630 | 671 | 327 |
| **SSC** | 183 | 184 | 198 | 210 | 100 |
| **IR** | 43 | 32 | 33 | 68 | 82 |

*Nicotiana tabacum* was used as reference for SNPs detection.
