## Supplementary material for "Plastid genomics of *Nicotiana* (Solanaceae): insights into molecular evolution, positive selection and the origin of the maternal genome of Aztec tobacco (*Nicotiana rustica*)": Table 3

**Table 3.** InDels distribution of *Nicotiana* chloroplast genome

|  | ***Nicotiana knightiana*** | **InDel length (bp)** | **InDel average length** |
| --- | --- | --- | --- |
| LSC | 91 | 506 | 5.56 |
| SSC | 11 | 36 | 3.27 |
| IR | 8 | 29 | 3.62 |
|  | ***Nicotiana rustica*** | **InDel length (bp)** | **InDel average length** |
| LSC | 89 | 478 | 5.37 |
| SSC | 11 | 36 | 3.27 |
| IR | 7 | 38 | 5.42 |
|  | ***Nicotiana paniculata*** | **InDel length (bp)** | **InDel average length** |
| LSC | 92 | 618 | 6.71 |
| SSC | 14 | 156 | 11.14 |
| IR | 10 | 28 | 2.80 |
|  | ***Nicotiana obtusifolia*** |  |  |
| LSC | 117 | 677 | 5.78 |
| SSC | 12 | 52 | 4.33 |
| IR | 14 | 167 | 11.92 |
|  | ***Nicotiana glauca*** | **InDel length (bp)** | **InDel average length** |
| LSC | 88 | 450 | 5.11 |
| SSC | 11 | 44 | 4 |
| IR | 14 | 82 | 5.85 |
