## Supplementary material for "Plastid genomics of *Nicotiana* (Solanaceae): insights into molecular evolution, positive selection and the origin of the maternal genome of Aztec tobacco (*Nicotiana rustica*)": Table 4

**Table 4.** Mutational hotspots among *Nicotiana* species

| **No** | **Region** | **Nucleotide Diversity** | **Total Number of Mutation** | **Region Length** |
| --- | --- | --- | --- | --- |
| 1 | *inf*A | 0.2594 | 45 | 249 |
| 2 | *rps*12 intron | 0.1527 | 161 | 527 |
| 3 | *rps*16*-trn*Q-UUG | 0.0845 | 225 | 1266 |
| 4 | *trn*K*-*UUU*-rps*16 | 0.048 | 46 | 703 |
| 5 | *trn*H*-psb*A | 0.0438 | 19 | 433 |
| 6 | *rpl*36*-inf*A | 0.0294 | 3 | 116 |
| 7 | *ccs*A*-ndh*D | 0.0287 | 17 | 237 |
| 8 | *rps*16*-*intron | 0.0278 | 27 | 862 |
| 9 | *rpl*32*-trn*L-UAG | 0.0261 | 61 | 931 |
| 10 | *trn*M*-*CAU*-atp*E | 0.0224 | 24 | 218 |
| 11 | *rpl*33*-rps*18 | 0.0222 | 20 | 180 |
| 12 | *pet*D*-rpo*A | 0.0198 | 9 | 182 |
| 13 | *rpl*14*-rpl*16 | 0.0184 | 10 | 119 |
| 14 | *ndh*E*-ndh*G | 0.0173 | 7 | 219 |
| 15 | *rps*15*-ycf*1 | 0.0171 | 17 | 385 |
| 16 | *ndh*H*-rps*15 | 0.0166 | 4 | 108 |
| 17 | *pet*G*-trn*W*-*CCA | 0.0157 | 4 | 127 |
| 18 | *pet*L*-pet*G | 0.0153 | 6 | 182 |
| 19 | *trn*G*-*GCC*-trnf*M | 0.0152 | 11 | 228 |
| 20 | *rpo*A*-rps*11 | 0.0151 | 2 | 66 |
