## Supplementary Table 1 for "Plastid genomics of *Nicotiana* (Solanaceae): insights into molecular evolution, positive selection and the origin of the maternal genome of Aztec tobacco (*Nicotiana rustica*)"

**Table 1S -** Base composition in the *Nicotiana knightiana* plastid genome

|  | **A%** | **C%** | **G%** | **T%** | **Length (bp)** |
| --- | --- | --- | --- | --- | --- |
| **Total** | 30.7 | 19.2 | 18.6 | 31.5 | 154,386 |
| **LSC** | 31.4 | 18.4 | 17.6 | 32.7 | 86,626 |
| **SSC** | 33.6 | 16.8 | 15.3 | 34.1 | 18,551 |
| **IR** | 26.3 | 22.4 | 20.7 | 28.5 | 25,364 |
| **tRNA** | 23.9 | 26.7 | 26.2 | 23.2 | 2790 |
| **rRNA** | 22.4 | 27.6 | 27.6 | 22.4 | 9052 |
| **Protein Coding genes** | 30.4 | 19.5 | 18.7 | 31.4 | 79,917 |
| **1st Position** | 30.9 | 19.0 | 18.7 | 31.4 | 51,990 |
| **2nd position** | 30.1 | 19.3 | 18.7 | 31.9 | 51,990 |
| **3rd position** | 31.1 | 19.4 | 18.5 | 31.1 | 51,989 |

**Table 1Sa -** Base composition in the *Nicotiana rustica* plastid genome

|  | **A%** | **C%** | **G%** | **T%** | **Length (bp)** |
| --- | --- | --- | --- | --- | --- |
| **Total** | 30.7 | 19.2 | 18.6 | 31.5 | 155,849 |
| **LSC** | 31.4 | 18.4 | 17.6 | 32.7 | 86,612 |
| **SSC** | 33.6 | 16.8 | 15.3 | 34.1 | 18,551 |
| **IR** | 28.3 | 22.4 | 20.8 | 28.5 | 25,342 |
| **tRNA** | 23.9 | 26.7 | 26.2 | 23.2 | 2790 |
| **rRNA** | 22.3 | 27.7 | 27.7 | 22.3 | 9050 |
| **Protein Coding genes** | 30.4 | 19.5 | 18.7 | 31.4 | 79,917 |
| **1st Position** | 30.3 | 19.1 | 18.5 | 31.4 | 51,950 |
| **2nd position** | 30.8 | 18.8 | 18.7 | 31.7 | 51,950 |
| **3rd position** | 30.3 | 19.8 | 18.6 | 31.3 | 51,949 |

**Table 1Sb -** Base composition in the *Nicotiana paniculata* plastid genome

|  | **A%** | **C%** | **G%** | **T%** | **Length (bp)** |
| --- | --- | --- | --- | --- | --- |
| **Total** | 30.7 | 19.2 | 18.6 | 31.5 | 155,689 |
| **LSC** | 31.4 | 18.4 | 17.6 | 32.7 | 86,510 |
| **SSC** | 33.7 | 16.8 | 15.4 | 34.1 | 18,441 |
| **IR** | 28.3 | 22.4 | 20.8 | 28.5 | 25,369 |
| **tRNA** | 23.9 | 26.7 | 26.2 | 23.2 | 2790 |
| **rRNA** | 22.3 | 27.7 | 27.7 | 22.3 | 9052 |
| **Protein Coding genes** | 30.4 | 19.5 | 18.7 | 31.4 | 79,896 |
| **1st Position** | 30.5 | 18.8 | 18.4 | 32.2 | 51,914 |
| **2nd position** | 29.7 | 19.9 | 19.5 | 30.9 | 51,913 |
| **3rd position** | 31.8 | 19.0 | 18.0 | 31.3 | 51,913 |

**Table 1Sc -** Base composition in the *Nicotiana obtusifolia* plastid genome

|  | **A%** | **C%** | **G%** | **T%** | **Length (bp)** |
| --- | --- | --- | --- | --- | --- |
| **Total** | 30.7 | 19.2 | 18.6 | 31.5 | 156,022 |
| **LSC** | 31.4 | 18.3 | 17.5 | 32.7 | 86,609 |
| **SSC** | 33.6 | 16.8 | 15.2 | 34.3 | 18,541 |
| **IR** | 28.3 | 22.4 | 20.8 | 28.5 | 25,436 |
| **tRNA** | 23.8 | 26.7 | 26.3 | 23.2 | 2797 |
| **rRNA** | 22.4 | 27.6 | 27.6 | 22.4 | 9050 |
| **Protein Coding genes** | 30.4 | 19.5 | 18.7 | 31.5 | 80,139 |
| **1st Position** | 31.1 | 19.2 | 18.5 | 31.2 | 52,008 |
| **2nd position** | 31.1 | 19.0 | 18.6 | 31.3 | 52,007 |
| **3rd position** | 29.9 | 19.4 | 18.7 | 32.0 | 52,007 |

**Table 1Sd -** Base composition in the *Nicotiana glauca* plastid genome

|  | **A%** | **C%** | **G%** | **T%** | **Length (bp)** |
| --- | --- | --- | --- | --- | --- |
| **Total** | 30.7 | 19.2 | 18.6 | 31.5 | 155,917 |
| **LSC** | 31.4 | 18.4 | 17.6 | 32.7 | 86,716 |
| **SSC** | 33.7 | 16.8 | 15.2 | 34.2 | 18,555 |
| **IR** | 28.4 | 22.4 | 20.8 | 28.5 | 25,323 |
| **tRNA** | 23.8 | 26.7 | 26.3 | 23.2 | 2797 |
| **rRNA** | 22.4 | 27.6 | 27.6 | 22.4 | 9052 |
| **Protein Coding genes** | 30.4 | 19.5 | 18.7 | 31.5 | 79,860 |
| **1st Position** | 31.5 | 18.6 | 18.4 | 31.5 | 51,973 |
| **2nd position** | 30.5 | 19.3 | 18.6 | 31.7 | 51,972 |
| **3rd position** | 30.1 | 19.9 | 18.8 | 31.3 | 51,972 |
