## Supplementary Table 2 for "Plastid genomics of *Nicotiana* (Solanaceae): insights into molecular evolution, positive selection and the origin of the maternal genome of Aztec tobacco (*Nicotiana rustica*)"

**Table S2** – RSCU (Relative synonymous codon usage) in the chloroplast genome of *N. knightiana, N. rustica, N. paniculata, N. obtusifolia and N. glauca*.

| **Codon** | **Amino acid** | **Number of codons** | | |  |  | **Codon** | **Amino acid** | **Number of Codons** | | |  |  |
| --- | --- | --- | --- | --- | --- | --- | --- | --- | --- | --- | --- | --- | --- |
|  |  | ***NK*** | ***NR*** | ***NP*** | ***NO*** | ***NG*** |  |  | ***NK*** | ***NR*** | ***NP*** | ***NO*** | ***NG*** |
| GCA | A | 405 | 405 | 401 | 407 | 404 | CCA | P | 341 | 347 | 333 | 331 | 331 |
| GCC | A | 253 | 253 | 248 | 244 | 247 | CCC | P | 223 | 225 | 216 | 213 | 213 |
| GCG | A | 142 | 144 | 139 | 136 | 136 | CCG | P | 166 | 169 | 159 | 154 | 157 |
| GCT | A | 628 | 626 | 625 | 630 | 628 | CCT | P | 444 | 443 | 429 | 438 | 434 |
| TGC | C | 85 | 88 | 80 | 78 | 79 | CAA | Q | 731 | 745 | 725 | 726 | 725 |
| TGT | C | 229 | 230 | 220 | 222 | 223 | CAG | Q | 223 | 246 | 241 | 244 | 242 |
| GAC | D | 226 | 229 | 220 | 215 | 221 | AGA | R | 509 | 523 | 486 | 493 | 487 |
| GAT | D | 911 | 925 | 880 | 884 | 875 | AGG | R | 190 | 191 | 175 | 174 | 179 |
| GAA | E | 1,088 | 1,092 | 1055 | 1056 | 1050 | CGA | R | 408 | 412 | 398 | 396 | 395 |
| GAG | E | 373 | 379 | 352 | 352 | 355 | CGC | R | 104 | 108 | 99 | 95 | 97 |
| TTC | F | 570 | 586 | 539 | 552 | 538 | CGG | R | 132 | 135 | 123 | 126 | 123 |
| TTT | F | 986 | 1010 | 967 | 970 | 966 | CGT | R | 350 | 352 | 347 | 341 | 342 |
| GGA | G | 756 | 763 | 731 | 738 | 731 | AGC | S | 127 | 128 | 121 | 120 | 120 |
| GGC | G | 205 | 206 | 204 | 208 | 204 | AGT | S | 430 | 429 | 411 | 413 | 410 |
| GGG | G | 324 | 329 | 318 | 317 | 322 | TCA | S | 432 | 440 | 411 | 412 | 407 |
| GGT | G | 594 | 595 | 574 | 570 | 574 | TCC | S | 355 | 360 | 339 | 338 | 336 |
| CAC | H | 152 | 152 | 147 | 148 | 149 | TCG | S | 216 | 220 | 197 | 204 | 205 |
| CAT | H | 502 | 510 | 482 | 485 | 485 | TCT | S | 620 | 634 | 603 | 601 | 602 |
| ATA | I | 723 | 744 | 695 | 694 | 694 | ACA | T | 428 | 426 | 409 | 418 | 415 |
| ATC | I | 472 | 482 | 457 | 455 | 458 | ACC | T | 275 | 275 | 267 | 271 | 270 |
| ATT | I | 1,122 | 1,134 | 1,100 | 1104 | 1101 | ACG | T | 153 | 153 | 153 | 148 | 148 |
| AAA | K | 1,084 | 1,121 | 1,052 | 1056 | 1052 | ACT | T | 546 | 547 | 526 | 525 | 524 |
| AAG | K | 388 | 395 | 386 | 379 | 382 | GTA | V | 563 | 564 | 536 | 536 | 542 |
| CTA | L | 395 | 408 | 375 | 374 | 373 | GTC | V | 206 | 206 | 185 | 187 | 185 |
| CTC | L | 231 | 231 | 213 | 211 | 213 | GTG | V | 216 | 216 | 202 | 203 | 199 |
| CTG | L | 200 | 198 | 196 | 197 | 194 | GTT | V | 545 | 551 | 532 | 534 | 529 |
| CTT | L | 642 | 647 | 620 | 619 | 629 | TGG | W | 492 | 495 | 486 | 483 | 485 |
| TTA | L | 895 | 905 | 883 | 883 | 880 | TAC | Y | 196 | 205 | 193 | 193 | 192 |
| TTG | L | 609 | 617 | 582 | 574 | 576 | TAT | Y | 792 | 797 | 775 | 778 | 774 |
| ATG | M | 662 | 679 | 630 | 625 | 630 | TAA | * | 51 | 57 | 47 | 46 | 46 |
| CTG | M | 1 | 2 | - | - | - |  |  |  |  |  |  |  |
| GTG | M | 5 | 4 | 4 | - | 4 |  |  |  |  |  |  |  |
| AAC | N | 326 | 337 | 319 | 314 | 315 | TAG | * | 22 | 28 | 24 | 23 | 24 |
| AAT | N | 1044 | 1064 | 997 | 1002 | 1001 | TGA | * | 21 | 22 | 17 | 19 | 18 |
