## Supplementary Table 3 for "Plastid genomics of *Nicotiana* (Solanaceae): insights into molecular evolution, positive selection and the origin of the maternal genome of Aztec tobacco (*Nicotiana rustica*)"

**Table S3**. Putative RNA editing sites in the *Nicotiana knightiana* chloroplast genome.

| **Gene** | **Nucleotide Position** | **Amino acid (AA) Position** | **Alignment Column** | **Effect** | **Score** |
| --- | --- | --- | --- | --- | --- |
| *atp*A | 791 | 264 | 264 | CCC (P) => CTC (L) | 1 |
| *atp*F | 92 | 31 | 31 | CCA (P) => CTA (L) | 0.86 |
| *ndh*A | 341 | 114 | 114 | TCA (S) => TTA (L) | 1 |
|  | 1073 | 358 | 358 | TCC (S) => TTC (F) | 1 |
| *ndh*B | 149 | 50 | 50 | TCA (S) => TTA (L) | 1 |
|  | 467 | 156 | 156 | CCA (P) => CTA (L) | 1 |
|  | 586 | 196 | 196 | CAT (H) => TAT (Y) | 1 |
|  | 611 | 204 | 204 | TCA (S) => TTA (L) | 0.8 |
|  | 737 | 246 | 246 | CCA (P) => CTA (L) | 1 |
|  | 746 | 249 | 249 | TCT (S) => TTT (F) | 1 |
|  | 830 | 277 | 277 | TCA (S) => TTA (L) | 1 |
|  | 836 | 279 | 279 | TCA (S) => TTA (L) | 1 |
|  | 1481 | 494 | 494 | CCA (P) => CTA (L) | 1 |
| *ndh*D | 383 | 128 | 128 | TCA (S) => TTA (L) | 1 |
|  | 599 | 200 | 200 | TCA (S) => TTA (L) | 1 |
|  | 674 | 225 | 225 | TCG (S) => TTG (L) | 1 |
|  | 878 | 293 | 293 | TCA (S) => TTA (L) | 1 |
|  | 1298 | 433 | 433 | TCA (S) => TTA (L) | 0.8 |
|  | 1310 | 437 | 437 | TCA (S) => TTA (L) | 0.8 |
| *ndh*F | 290 | 97 | 97 | TCA (S) => TTA (L) | 1 |
| *pet*B | 614 | 205 | 205 | CCA (P) => CTA (L) | 1 |
| *psb*E | 214 | 72 | 72 | CCT (P) => TCT (S) | 1 |
| *rpl*20 | 308 | 103 | 103 | TCA (S) => TTA (L) | 0.86 |
| rpoA | 830 | 277 | 279 | TCA (S) => TTA (L) | 1 |
| *rpo*B | 338 | 113 | 113 | TCT (S) => TTT (F) | 1 |
|  | 473 | 158 | 159 | TCA (S) => TTA (L) | 0.86 |
|  | 551 | 184 | 185 | TCA (S) => TTA (L) | 1 |
|  | 2000 | 667 | 684 | TCT (S) => TTT (F) | 1 |
| *rpo*C1 | 62 | 21 | 21 | TCA (S) => TTA (L) | 1 |
| *rpo*C2 | 2287 | 763 | 948 | CGG (R) => TGG (W) | 1 |
|  | 3731 | 1244 | 1450 | TCA (S) => TTA (L) | 0.86 |
| *rps*2 | 248 | 83 | 83 | TCA (S) => TTA (L) | 1 |
| *rps*14 | 80 | 27 | 27 | TCA (S) => TTA (L) | 1 |
|  | 149 | 50 | 53 | CCA (P) => CTA (L) | 1 |

**Table S3a**. Putative RNA editing sites in the *Nicotiana rustica* chloroplast genome.

| **Gene** | **Nucleotide Position** | **Amino acid (AA) Position** | **Alignment Column** | **Effect** | **Score** |
| --- | --- | --- | --- | --- | --- |
| *atp*A | 791 | 264 | 264 | CCC (P) => CTC (L) | 1 |
| *atp*F | 92 | 31 | 31 | CCA (P) => CTA (L) | 0.86 |
| *ndh*A | 341 | 114 | 114 | TCA (S) => TTA (L) | 1 |
|  | 1073 | 358 | 358 | TCC (S) => TTC (F) | 1 |
| *ndh*B | 149 | 50 | 50 | TCA (S) => TTA (L) | 1 |
|  | 467 | 156 | 156 | CCA (P) => CTA (L) | 1 |
|  | 586 | 196 | 196 | CAT (H) => TAT (Y) | 1 |
|  | 611 | 204 | 204 | TCA (S) => TTA (L) | 0.8 |
|  | 737 | 246 | 246 | CCA (P) => CTA (L) | 1 |
|  | 746 | 249 | 249 | TCT (S) => TTT (F) | 1 |
|  | 830 | 277 | 277 | TCA (S) => TTA (L) | 1 |
|  | 836 | 279 | 279 | TCA (S) => TTA (L) | 1 |
|  | 1481 | 494 | 494 | CCA (P) => CTA (L) | 1 |
| *ndh*D | 383 | 128 | 128 | TCA (S) => TTA (L) | 1 |
|  | 599 | 200 | 200 | TCA (S) => TTA (L) | 1 |
|  | 674 | 225 | 225 | TCG (S) => TTG (L) | 1 |
|  | 878 | 293 | 293 | TCA (S) => TTA (L) | 1 |
|  | 1298 | 433 | 433 | TCA (S) => TTA (L) | 0.8 |
|  | 1310 | 437 | 437 | TCA (S) => TTA (L) | 0.8 |
| *ndh*F | 290 | 97 | 97 | TCA (S) => TTA (L) | 1 |
| *pet*B | 611 | 204 | 204 | CCA (P) => CTA (L) | 1 |
| *psb*E | 214 | 72 | 72 | CCT (P) => TCT (S) | 1 |
| *rpl*20 | 308 | 103 | 103 | TCA (S) => TTA (L) | 0.86 |
| *rpo*A | 830 | 277 | 279 | TCA (S) => TTA (L) | 1 |
| *rpo*B | 338 | 113 | 113 | TCT (S) => TTT (F) | 1 |
|  | 473 | 158 | 159 | TCA (S) => TTA (L) | 0.86 |
|  | 551 | 184 | 185 | TCA (S) => TTA (L) | 1 |
|  | 2000 | 667 | 684 | TCT (S) => TTT (F) | 1 |
| *rpo*C1 | 62 | 21 | 21 | TCA (S) => TTA (L) | 1 |
|  | 2287 | 763 | 948 | CGG (R) => TGG (W) |  |
| *rpo*C2 | 3731 | 1244 | 1450 | TCA (S) => TTA (L) | 0.86 |
| rps2 | 248 | 83 | 83 | TCA (S) => TTA (L) | 1 |
| *rps*14 | 80 | 27 | 27 | TCA (S) => TTA (L) | 1 |
|  | 149 | 50 | 53 | CCA (P) => CTA (L) | 1 |

**Table S3b** **-** Putative RNA editing sites in the *Nicotiana paniculata* chloroplast genome.

| **Gene** | **Nucleotide Position** | **Amino acid (AA) Position** | **Alignment Column** | **Effect** | **Score** |
| --- | --- | --- | --- | --- | --- |
| *atp*A | 791 | 264 | 264 | CCC (P) => CTC (L) | 1 |
| *atp*F | 92 | 31 | 31 | CCA (P) => CTA (L) | 0.86 |
| *ndh*A | 341 | 114 | 114 | TCA (S) => TTA (L) |  |
|  | 1073 | 358 | 358 | TCC (S) => TTC (F) | 1 |
| *ndh*B | 149 | 50 | 50 | TCA (S) => TTA (L) | 1 |
|  | 467 | 156 | 156 | CCA (P) => CTA (L) | 1 |
|  | 586 | 196 | 196 | CAT (H) => TAT (Y) | 1 |
|  | 611 | 204 | 204 | TCA (S) => TTA (L) | 0.8 |
|  | 737 | 246 | 246 | CCA (P) => CTA (L) | 1 |
|  | 746 | 249 | 249 | TCT (S) => TTT (F) | 1 |
|  | 830 | 277 | 277 | TCA (S) => TTA (L) | 1 |
|  | 836 | 279 | 279 | TCA (S) => TTA (L) | 1 |
|  | 1481 | 494 | 494 | CCA (P) => CTA (L) | 1 |
| *ndh*D | 383 | 128 | 128 | TCA (S) => TTA (L) | 1 |
|  | 599 | 200 | 200 | TCA (S) => TTA (L) | 1 |
|  | 674 | 225 | 225 | TCG (S) => TTG (L) | 1 |
|  | 878 | 293 | 293 | TCA (S) => TTA (L) | 1 |
|  | 1298 | 433 | 433 | TCA (S) => TTA (L) | 0.8 |
|  | 1310 | 437 | 437 | TCA (S) => TTA (L) | 0.8 |
| *ndh*F | 290 | 97 | 97 | TCA (S) => TTA (L) | 1 |
| *pet*B | 611 | 204 | 204 | CCA (P) => CTA (L) | 1 |
| *psb*E | 214 | 72 | 72 | CCT (P) => TCT (S) | 1 |
| *rpl*20 | 308 | 103 | 103 | TCA (S) => TTA (L) | 0.86 |
| *rpo*A | 830 | 277 | 279 | TCA (S) => TTA (L) | 1 |
| *rpo*B | 338 | 113 | 113 | TCT (S) => TTT (F) | 1 |
|  | 473 | 158 | 159 | TCA (S) => TTA (L) | 0.86 |
|  | 551 | 184 | 185 | TCA (S) => TTA (L) | 1 |
|  | 2000 | 667 | 684 | TCT (S) => TTT (F) | 1 |
| *rpo*C1 | 62 | 21 | 21 | TCA (S) => TTA (L) | 1 |
| *rpo*C2 | 2287 | 763 | 948 | CGG (R) => TGG (W) | 1 |
|  | 3731 | 1244 | 1450 | TCA (S) => TTA (L) | 0.86 |
| *rps*2 | 248 | 83 | 83 | TCA (S) => TTA (L) | 1 |
| *rps*14 | 80 | 27 | 27 | TCA (S) => TTA (L) | 1 |
|  | 149 | 50 | 53 | CCA (P) => CTA (L) | 1 |

**Table S3c** **-** Putative RNA editing sites in the *Nicotiana obtusifolia* chloroplast genome.

| **Gene** | **Nucleotide Position** | **Amino acid (AA) Position** | **Alignment Column** | **Effect** | **Score** |
| --- | --- | --- | --- | --- | --- |
| *atp*A | 791 | 264 | 264 | CCC (P) => CTC (L) | 1 |
| atpF | 92 | 31 | 31 | CCA (P) => CTA (L) | 0.86 |
| ndhA | 341 | 114 | 114 | TCA (S) => TTA (L) | 1 |
|  | 1073 | 358 | 358 | TCC (S) => TTC (F) | 1 |
| *ndh*B | 149 | 50 | 50 | TCA (S) => TTA (L) | 1 |
|  | 467 | 156 | 156 | CCA (P) => CTA (L) | 1 |
|  | 586 | 196 | 196 | CAT (H) => TAT (Y) | 1 |
|  | 611 | 204 | 204 | TCA (S) => TTA (L) | 0.8 |
|  | 737 | 246 | 246 | CCA (P) => CTA (L) | 1 |
|  | 746 | 249 | 249 | TCT (S) => TTT (F) | 1 |
|  | 830 | 277 | 277 | TCA (S) => TTA (L) | 1 |
|  | 836 | 279 | 279 | TCA (S) => TTA (L) | 1 |
|  | 1481 | 494 | 494 | CCA (P) => CTA (L) | 1 |
| *ndh*D | 29 | 10 | 10 | ACG (T) => ATG (M) | 1 |
|  | 410 | 137 | 137 | TCA (S) => TTA (L) | 1 |
|  | 626 | 209 | 209 | TCA (S) => TTA (L) | 1 |
|  | 701 | 234 | 234 | TCG (S) => TTG (L) | 1 |
|  | 905 | 302 | 302 | TCA (S) => TTA (L) | 1 |
|  | 1325 | 442 | 442 | TCA (S) => TTA (L) | 0.8 |
|  | 1337 | 446 | 446 | TCA (S) => TTA (L) | 0.8 |
| *ndh*F | 290 | 97 | 97 | TCA (S) => TTA (L) | 1 |
| *pet*B | 611 | 204 | 204 | CCA (P) => CTA (L) | 1 |
| *psb*E | 214 | 72 | 72 | CCT (P) => TCT (S) | 1 |
| *rpl*20 | 308 | 103 | 103 | TCA (S) => TTA (L) | 0.86 |
| *rpo*A | 830 | 277 | 279 | TCA (S) => TTA (L) | 1 |
| *rpo*B | 338 | 113 | 113 | TCT (S) => TTT (F) | 1 |
|  | 473 | 158 | 159 | TCA (S) => TTA (L) | 0.86 |
|  | 551 | 184 | 185 | TCA (S) => TTA (L) | 1 |
|  | 2000 | 667 | 684 | TCT (S) => TTT (F) | 1 |
| *rpo*C1 | 62 | 21 | 21 | TCA (S) => TTA (L) | 1 |
| *rpo*C2 | 2287 | 763 | 948 | CGG (R) => TGG (W) | 1 |
|  | 3731 | 1244 | 1450 | TCA (S) => TTA (L) | 0.86 |
| *rps*2 | 248 | 83 | 83 | TCA (S) => TTA (L) | 1 |
| *rps*14 | 80 | 27 | 27 | TCA (S) => TTA (L) | 1 |
|  | 149 | 50 | 53 | CCA (P) => CTA (L) | 1 |

**Table S3d** - Putative RNA editing sites in the *Nicotiana glauca* chloroplast genome.

| **Gene** | **Nucleotide**  **Position** | **Amino acid**  **(AA)**  **Position** | **Alignment**  **Column** | **Effect** | **Score** |
| --- | --- | --- | --- | --- | --- |
| *atp*A | 791 | 264 | 264 | CCC (P) => CTC (L) | 1 |
| *atp*F | 92 | 31 | 31 | CCA (P) => CTA (L) | 0.86 |
| *ndh*A | 341 | 114 | 114 | TCA (S) => TTA (L) | 1 |
|  | 1073 | 358 | 358 | TCC (S) => TTC (F) | 1 |
| *ndh*B | 149 | 50 | 50 | TCA (S) => TTA (L) | 1 |
|  | 467 | 156 | 156 | CCA (P) => CTA (L) | 1 |
|  | 586 | 196 | 196 | CAT (H) => TAT (Y) | 1 |
|  | 611 | 204 | 204 | TCA (S) => TTA (L) | 0.8 |
|  | 737 | 246 | 246 | CCA (P) => CTA (L) | 1 |
|  | 746 | 249 | 249 | TCT (S) => TTT (F) | 1 |
|  | 830 | 277 | 277 | TCA (S) => TTA (L) | 1 |
|  | 836 | 279 | 279 | TCA (S) => TTA (L) | 1 |
|  | 1481 | 494 | 494 | CCA (P) => CTA (L) | 1 |
| *ndh*D | 410 | 137 | 137 | TCA (S) => TTA (L) | 1 |
|  | 626 | 209 | 209 | TCA (S) => TTA (L) | 1 |
|  | 701 | 234 | 234 | TCG (S) => TTG (L) | 1 |
|  | 853 | 285 | 285 | CTT (L) => TTT (F) | 1 |
|  | 905 | 302 | 302 | TCA (S) => TTA (L) | 1 |
|  | 1325 | 442 | 442 | TCA (S) => TTA (L) | 0.8 |
|  | 1337 | 446 | 446 | TCA (S) => TTA (L) | 0.8 |
|  | 410 | 137 | 137 | TCA (S) => TTA (L) | 1 |
| *ndh*F | 290 | 97 | 97 | TCA (S) => TTA (L) | 1 |
| *pet*B | 611 | 204 | 204 | CCA (P) => CTA (L) | 1 |
| *psb*E | 214 | 72 | 72 | CCT (P) => TCT (S) | 1 |
| *rpl*20 | 308 | 103 | 103 | TCA (S) => TTA (L) | 0.86 |
| *rpo*A | 830 | 277 | 279 | TCA (S) => TTA (L) | 1 |
| *rpo*B | 338 | 113 | 113 | TCT (S) => TTT (F) | 1 |
|  | 473 | 158 | 159 | TCA (S) => TTA (L) | 0.86 |
|  | 551 | 184 | 185 | TCA (S) => TTA (L) | 1 |
|  | 2000 | 667 | 684 | TCT (S) => TTT (F) | 1 |
|  | 338 | 113 | 113 | TCT (S) => TTT (F) | 1 |
| *rpo*C1 | 62 | 21 | 21 | TCA (S) => TTA (L) | 1 |
| *rpo*C2 | 2287 | 763 | 948 | CGG (R) => TGG (W) | 1 |
|  | 3731 | 1244 | 1450 | TCA (S) => TTA (L) | 0.86 |
| *rps*2 | 248 | 83 | 83 | TCA (S) => TTA (L) | 1 |
| *rps*14 | 80 | 27 | 27 | TCA (S) => TTA (L) | 1 |
|  | 149 | 50 | 53 | CCA (P) => CTA (L) | 1 |
