## Supplementary Table 4 for "Plastid genomics of *Nicotiana* (Solanaceae): insights into molecular evolution, positive selection and the origin of the maternal genome of Aztec tobacco (*Nicotiana rustica*)"

**Table S4 -** Comparison of synonymous and non-synonymous substitution in *Nicotiana* species

|  |  | Ks | Ka | Ka/Ks |
| --- | --- | --- | --- | --- |
| *psb*A | NG | 0 | 0 | 0 |
|  | NK | 0 | 0.0048 | 0 |
|  | NO | 0.0057 | 0 | 0 |
|  | NP | 0 | 0.0048 | 0 |
|  | NR | 0 | 0.0047 | 0 |
| *rps*16 | NG | 0 | 0 | 0 |
|  | NK | 0 | 0 | 0 |
|  | NO | 0 | 0 | 0 |
|  | NP | 0 | 0 | 0 |
|  | NR | 0 | 0 | 0 |
| *psb*K | NG | 0 | 0 | 0 |
|  | NK | 0.0509 | 0 | 0 |
|  | NO | 0.025 | 0 | 0 |
|  | NP | 0 | 0.0146 | 0 |
|  | NR | 0.0473 | 0 | 0 |
| *psb*I | NG | 0 | 0.0109 | 0 |
|  | NK | 0 | 0.0109 | 0 |
|  | NO | 0 | 0.0109 | 0 |
|  | NP | 0 | 0.022 | 0 |
|  | NR | 0 | 0.0107 | 0 |
| *atp*A | NG | 0 | 0.0025 | 0 |
|  | NK | 0.0094 | 0.0059 | 0.6276 |
|  | NO | 0 | 0.005 | 0 |
|  | NP | 0.0094 | 0.0067 | 0.71276 |
|  | NR | 0.0094 | 0.0058 | 0.61702 |
| *atp*F | NG | 0 | 0 | 0 |
|  | NK | 0 | 0.0062 | 0 |
|  | NO | 0.0124 | 0.0156 | 1.2580 |
|  | NP | 0 | 0.0062 | 0 |
|  | NR | 0 | 0.0062 | 0 |
| *atp*H | NG | 0 | 0 | 0 |
|  | NK | 0 | 0.0104 | 0 |
|  | NO | 0 | 0.0104 | 0 |
|  | NP | 0 | 0.0052 | 0 |
|  | NR | 0 | 0.0051 | 0 |
| *atp*I | NG | 0 | 0.0017 | 0 |
|  | NK | 0 | 0.0033 | 0 |
|  | NO | 0 | 0 | 0 |
|  | NP | 0.0027 | 0 | 0 |
|  | NR | 0 | 0.0033 | 0 |
| *rps*2 | NG | 0 | 0.0018 | 0 |
|  | NK | 0.0069 | 0.0072 | 1.0434 |
|  | NO | 0 | 0.0054 | 0 |
|  | NP | 0.0069 | 0.0072 | 1.0434 |
|  | NR | 0.0068 | 0.0071 | 1.0441 |
| *rpo*C2 | NG | 0.0035 | 0.0027 | 0.7714 |
|  | NK | 0.0058 | 0.004 | 0.6896 |
|  | NO | 0.0069 | 0.0049 | 0.7101 |
|  | NP | 0.0058 | 0.0037 | 0.6379 |
|  | NR | 0.0058 | 0.004 | 0.6896 |
| *rpo*C1 | NG | 0 | 0.0024 | 0 |
|  | NK | 0.003 | 0.0047 | 1.5666 |
|  | NO | 0.006 | 0.0047 | 0.7833 |
|  | NP | 0.003 | 0.0039 | 1.3 |
|  | NR | 0.003 | 0.0047 | 1.5666 |
| *rpo*B | NG | 0 | 0.0012 | 0 |
|  | NK | 0.003 | 0.0024 | 0.8 |
|  | NO | 0 | 0.0043 | 0 |
|  | NP | 0.003 | 0.0024 | 0.8 |
|  | NR | 0.003 | 0.0024 | 0.8 |
| *pet*N | NG | 0 | 0 | 0 |
|  | NK | 0 | 0 | 0 |
|  | NO | 0 | 0 | 0 |
|  | NP | 0 | 0 | 0 |
|  | NR | 0 | 0 | 0 |
| *psb*M | NG | 0 | 0 | 0 |
|  | NK | 0 | 0 | 0 |
|  | NO | 0 | 0 | 0 |
|  | NP | 0 | 0 | 0 |
|  | NR | 0 | 0 | 0 |
| *psb*D | NG | 0.004 | 0 | 0 |
|  | NK | 0.0163 | 0 | 0 |
|  | NO | 0.0161 | 0.0012 | 0.0745 |
|  | NP | 0.0161 | 0 | 0 |
|  | NR | 0.0161 | 0 | 0 |
| *psb*C | NG | 0.0059 | 0 | 0 |
|  | NK | 0.0178 | 0 | 0 |
|  | NO | 0.0089 | 0.0019 | 0.2134 |
|  | NP | 0.0178 | 0 | 0 |
|  | NR | 0.0177 | 0 | 0 |
| *psb*Z | NG | 0 | 0 | 0 |
|  | NK | 0.0219 | 0 | 0 |
|  | NO | 0.0219 | 0 | 0 |
|  | NP | 0.0219 | 0 | 0 |
|  | NR | 0 | 0 | 0 |
| *rps*14 | NG | 0 | 0 | 0 |
|  | NK | 0 | 0.0043 | 0 |
|  | NO | 0.0146 | 0.0043 | 0.2945 |
|  | NP | 0 | 0.0043 | 0 |
|  | NR | 0 | 0.0043 | 0 |
| *psa*B | NG | 0 | 0.0006 | 0 |
|  | NK | 0 | 0.004 | 0 |
|  | NO | 0 | 0.0045 | 0 |
|  | NP | 0 | 0.0034 | 0 |
|  | NR | 0 | 0.0034 | 0 |
| *psa*A | NG | 0.0022 | 0.0017 | 0.7727 |
|  | NK | 0 | 0.0022 | 0 |
|  | NO | 0 | 0.0017 | 0 |
|  | NP | 0 | 0.0022 | 0 |
|  | NR | 0 | 0.0022 | 0 |
| *ycf*3 | NG | 0 | 0 | 0 |
|  | NK | 0 | 0 | 0 |
|  | NO | 0 | 0 | 0 |
|  | NP | 0 | 0 | 0 |
|  | NR | 0 | 0 | 0 |
| *rps*4 | NG | 0 | 0.0021 | 0 |
|  | NK | 0 | 0.0021 | 0 |
|  | NO | 0 | 0.0063 | 0 |
|  | NP | 0 | 0.0021 | 0 |
|  | NR | 0 | 0.0021 | 0 |
| *ndh*J | NG | 0 | 0 | 0 |
|  | NK | 0 | 0.0027 | 0 |
|  | NO | 0 | 0 | 0 |
|  | NP | 0 | 0.0027 | 0 |
|  | NR | 0 | 0.0027 | 0 |
| *ndh*K | NG | 0.0067 | 0.0017 | 0.2537 |
|  | NK | 0.0067 | 0.0034 | 0.5074 |
|  | NO | 0.0134 | 0.0034 | 0.2537 |
|  | NP | 0.0067 | 0.0034 | 0.5074 |
|  | NR | 0.0066 | 0.0034 | 0.5151 |
| *ndh*C | NG | 0 | 0 | 0 |
|  | NK | 0 | 0.0068 | 0 |
|  | NO | 0.0145 | 0.0068 | 0.4689 |
|  | NP | 0 | 0.0068 | 0 |
|  | NR | 0 | 0.0068 | 0 |
| *atp*E | NG | 0 | 0.0032 | 0 |
|  | NK | 0.0025 | 0 | 0 |
|  | NO | 0.0123 | 0 | 0 |
|  | NP | 0.0123 | 0 | 0 |
|  | NR | 0.0122 | 0 | 0 |
| *atp*B | NG | 0 | 0.0043 | 0 |
|  | NK | 0.0032 | 0.0077 | 2.4062 |
|  | NO | 0.0032 | 0.0068 | 2.1250 |
|  | NP | 0.0032 | 0.0068 | 2.1250 |
|  | NR | 0.0032 | 0.0077 | 2.4062 |
| *rbc*L | NG | 0.0029 | 0.0009 | 0.3103 |
|  | NK | 0 | 0.0009 | 0 |
|  | NO | 0.0058 | 0.0028 | 0.4827 |
|  | NP | 0 | 0.0009 | 0 |
|  | NR | 0 | 0.0009 | 0 |
| *acc*D | NG | 0.0093 | 0.0033 | 0.3548 |
|  | NK | 0.6174 | 0.7088 | 1.1480 |
|  | NO | 0.0156 | 0.0083 | 0.5320 |
|  | NP | 0.0093 | 0.005 | 0.5376 |
|  | NR | 0.0092 | 0.0041 | 0.4456 |
| *psa*I | NG | 0 | 0 | 0 |
|  | NK | 0 | 0 | 0 |
|  | NO | 0 | 0 | 0 |
|  | NP | 0 | 0 | 0 |
|  | NR | 0 | 0 | 0 |
| *ycf*4 | NG | 0 | 0.0024 | 0 |
|  | NK | 0 | 0.0047 | 0 |
|  | NO | 0.0078 | 0.0047 | 0.6025 |
|  | NP | 0 | 0.0047 | 0 |
|  | NR | 0 | 0.0047 | 0 |
| *cem*A | NG | 0.0139 | 0.0018 | 0.1294 |
|  | NK | 0.028 | 0.0018 | 0.0642 |
|  | NO | 0.0279 | 0.0019 | 0.0681 |
|  | NP | 0 | 0.0093 | 0 |
|  | NR | 0.0274 | 0.0018 | 0.0656 |
| *pet*A | NG | 0.0044 | 0.0014 | 0.3181 |
|  | NK | 0.0088 | 0.0014 | 0.1590 |
|  | NO | 0.0268 | 0 | 0 |
|  | NP | 0.0088 | 0.0027 | 0.3068 |
|  | NR | 0.0087 | 0.0014 | 0.1609 |
| *psb*J | NG | 0 | 0.0107 | 0 |
|  | NK | 0.04 | 0.0215 | 0 |
|  | NO | 0 | 0.0107 | 0 |
|  | NP | 0.04 | 0.0215 | 0.5375 |
|  | NR | 0.038 | 0.0211 | 0.5552 |
| *psb*L | NG | 0 | 0.0109 | 0 |
|  | NK | 0 | 0.0109 | 0 |
|  | NO | 0 | 0.0109 | 0 |
|  | NP | 0 | 0.0220 | 0 |
|  | NR | 0 | 0.0107 | 0 |
| *psb*F | NG | 0 | 0 | 0 |
|  | NK | 0 | 0 | 0 |
|  | NO | 0 | 0.0107 | 0 |
|  | NP | 0 | 0 | 0 |
|  | NR | 0 | 0 | 0 |
| *psb*E | NG | 0 | 0 | 0 |
|  | NK | 0 | 0 | 0 |
|  | NO | 0 | 0 | 0 |
|  | NP | 0 | 0 | 0 |
|  | NR | 0 | 0 | 0 |
| *pet*L | NG | 0 | 0 | 0 |
|  | NK | 0 | 0 | 0 |
|  | NO | 0 | 0 | 0 |
|  | NP | 0 | 0 | 0 |
|  | NR | 0 | 0 | 0 |
| *pet*G | NG | 0 | 0 | 0 |
|  | NK | 0 | 0 | 0 |
|  | NO | 0 | 0 | 0 |
|  | NP | 0 | 0 | 0 |
|  | NR | 0 | 0 | 0 |
| *psa*J | NG | 0 | 0 | 0 |
|  | NK | 0.0305 | 0 | 0 |
|  | NO | 0 | 0 | 0 |
|  | NP | 0.0305 | 0 | 0 |
|  | NR | 0.0279 | 0 | 0 |
| *rpl*33 | NG | 0 | 0 | 0 |
|  | NK | 0 | 0 | 0 |
|  | NO | 0 | 0 | 0 |
|  | NP | 0 | 0 | 0 |
|  | NR | 0 | 0 | 0 |
| *rps*18 | NG | 0 | 0.0043 | 0 |
|  | NK | 0 | 0.0043 | 0 |
|  | NO | 0 | 0 | 0 |
|  | NP | 0 | 0.0043 | 0 |
|  | NR | 0 | 0.0043 | 0 |
| *rpl*20 | NG | 0 | 0 | 0 |
|  | NK | 0 | 0 | 0 |
|  | NO | 0 | 0.0034 | 0 |
|  | NP | 0 | 0 | 0 |
|  | NR | 0 | 0 | 0 |
| *rps*12 | NG | 0.0191 | 0.0056 | 0.2931 |
|  | NK | 0.0082 | 0.0089 | 1.0853 |
|  | NO | 0.0164 | 0 | 0 |
|  | NP | 0.0082 | 0.0089 | 1.0853 |
|  | NR | 0 | 0 | 0 |
| *clp*P | NG | 0 | 0 | 0 |
|  | NK | 0 | 0.0043 | 0 |
|  | NO | 0 | 0 | 0 |
|  | NP | 0 | 0.0043 | 0 |
|  | NR | 0 | 0.0043 | 0 |
| *psb*B | NG | 0.124 | 0.173 | 1.3951 |
|  | NK | 0.0195 | 0 | 0 |
|  | NO | 0.0167 | 0.0017 | 0.1017 |
|  | NP | 0.0195 | 0 | 0 |
|  | NR | 0.0193 | 0 | 0 |
| *psb*T | NG | 0 | 0 | 0 |
|  | NK | 0 | 0 | 0 |
|  | NO | 0 | 0 | 0 |
|  | NP | 0 | 0 | 0 |
|  | NR | 0 | 0 | 0 |
| *psb*N | NG | 0 | 0 | 0 |
|  | NK | 0 | 0 | 0 |
|  | NO | 0 | 0 | 0 |
|  | NP | 0 | 0 | 0 |
|  | NR | 0 | 0 | 0 |
| *psb*H | NG | 0 | 0 | 0 |
|  | NK | 0.0186 | 0 | 0 |
|  | NO | 0 | 0 | 0 |
|  | NP | 0.0186 | 0 | 0 |
|  | NR | 0.0045 | 0 | 0 |
| *pet*B | NG | 0.0064 | 0 | 0 |
|  | NK | 0.0064 | 0 | 0 |
|  | NO | 0.0064 | 0 | 0 |
|  | NP | 0.0064 | 0 | 0 |
|  | NR | 0.0063 | 0 | 0 |
| *pet*D | NG | 0 | 0 | 0 |
|  | NK | 0 | 0.0027 | 0 |
|  | NO | 0 | 0 | 0 |
|  | NP | 0 | 0.0027 | 0 |
|  | NR | 0 | 0.0026 | 0 |
| *rpo*A | NG | 0.0099 | 0.0037 | 0.3737 |
|  | NK | 0.0049 | 0.0112 | 2.2857 |
|  | NO | 0.0099 | 0.0062 | 0.6262 |
|  | NP | 0.0049 | 0.0112 | 2.2857 |
|  | NR | 0.0049 | 0.0112 | 2.2857 |
| *rps*11 | NG | 0 | 0 | 0 |
|  | NK | 0 | 0 | 0 |
|  | NO | 0.0106 | 0.0031 | 0.2924 |
|  | NP | 0 | 0 | 0 |
|  | NR | 0 | 0 | 0 |
| *rpl*36 | NG | 0 | 0 | 0 |
|  | NK | 0 | 0 | 0 |
|  | NO | 0 | 0 | 0 |
|  | NP | 0 | 0 | 0 |
|  | NR | 0 | 0 | 0 |
| *inf*A | NG | 3.8294 | 1.9972 | 0.5215 |
|  | NK | 0 | 0 | 0 |
|  | NO | 2.5509 | 2.7282 | 1.0695 |
|  | NP | 5.2691 | 2.3208 | 0.4404 |
|  | NR | 0 | 0 | 0 |
| *rps*8 | NG | 0 | 0.0032 | 0 |
|  | NK | 0 | 0.0032 | 0 |
|  | NO | 0 | 0.0064 | 0 |
|  | NP | 0 | 0.0032 | 0 |
|  | NR | 0 | 0.0032 | 0 |
| *rpl*14 | NG | 0 | 0 | 0 |
|  | NK | 0 | 0.0069 | 0 |
|  | NO | 0 | 0 | 0 |
|  | NP | 0 | 0.0069 | 0 |
|  | NR | 0 | 0.0068 | 0 |
| *rpl*16 | NG | 0.0101 | 0 | 0 |
|  | NK | 0 | 0.0066 | 0 |
|  | NO | 0 | 0.0033 | 0 |
|  | NP | 0 | 0.0066 | 0 |
|  | NR | 0 | 0.0065 | 0 |
| *rps*3 | NG | 0.0075 | 0.0039 | 0.52 |
|  | NK | 0 | 0.0097 | 0 |
|  | NO | 0.0075 | 0.0039 | 0.52 |
|  | NP | 0 | 0.0077 | 0 |
|  | NR | 0 | 0.0077 | 0 |
| *rpl*22 | NG | 0.0205 | 0.0027 | 0.1317 |
|  | NK | 0.0309 | 0.0054 | 0.1747 |
|  | NO | 0.0205 | 0.0109 | 0.5317 |
|  | NP | 0.0309 | 0.0054 | 0.1747 |
|  | NR | 0.0309 | 0.0054 | 0.1747 |
| *rps*19 | NG | 0 | 0.0091 | 0 |
|  | NK | 0 | 0.0091 | 0 |
|  | NO | 0 | 0.0091 | 0 |
|  | NP | 0.0073 | 0 | 0 |
|  | NR | 0 | 0.009 | 0 |
| *rpl*2 | NG | 0 | 0 | 0 |
|  | NK | 0 | 0 | 0 |
|  | NO | 0 | 0 | 0 |
|  | NP | 0 | 0 | 0 |
|  | NR | 0 | 0 | 0 |
| *rpl*23 | NG | 0 | 0 | 0 |
|  | NK | 0 | 0 | 0 |
|  | NO | 0 | 0 | 0 |
|  | NP | 0 | 0 | 0 |
|  | NR | 0 | 0 | 0 |
| *ycf*2 | NG | 0.7481 | 0.7235 | 0.9671 |
|  | NK | 0.0028 | 0.0015 | 0.5357 |
|  | NO | 0.749 | 0.7247 | 0.9675 |
|  | NP | 0.0159 | 0.0127 | 0.7987 |
|  | NR | 0.0028 | 0.0015 | 0.5357 |
| *ndh*B | NG | 0 | 0 | 0 |
|  | NK | 0.0047 | 0.0078 | 1.6595 |
|  | NO | 0 | 0 | 0 |
|  | NP | 0 | 0 | 0 |
|  | NR | 0 | 0 | 0 |
| *rps*7 | NG | 0 | 0 | 0 |
|  | NK | 0 | 0 | 0 |
|  | NO | 0 | 0 | 0 |
|  | NP | 0 | 0 | 0 |
|  | NR | 0 | 0 | 0 |
| *ycf*1 | NG | 1.2614 | 1.1461 | 0.9085 |
|  | NK | 0.0086 | 0.0051 | 0.5930 |
|  | NO | 0.0086 | 0.0051 | 0.5930 |
|  | NP | 0.0344 | 0.0285 | 0.8284 |
|  | NR | 0.0086 | 0.0051 | 0.5930 |
| *ndh*F | NG | 0.0024 | 0.0045 | 1.8750 |
|  | NK | 0.0047 | 0.0078 | 1.6595 |
|  | NO | 0.7625 | 0.73 | 0.9573 |
|  | NP | 0.0059 | 0.0098 | 1.6610 |
|  | NR | 0.0047 | 0.0078 | 1.6595 |
| *rpl*32 | NG | 0.0556 | 0 | 0 |
|  | NK | 0.0556 | 0 | 0 |
|  | NO | 0.0556 | 0 | 0 |
|  | NP | 0.3062 | 0.2116 | 0.6910 |
|  | NR | 0.0513 | 0 | 0 |
| *ccs*A | NG | 0 | 0 | 0 |
|  | NK | 0.0187 | 0.0028 | 0.1497 |
|  | NO | 0.0046 | 0.0042 | 0.9130 |
|  | NP | 0 | 0 | 0 |
|  | NR | 0.0184 | 0.0028 | 0.1521 |
| *ndh*D | NG | 0.0106 | 0.0033 | 0.3113 |
|  | NK | 0.0035 | 0.005 | 1.4285 |
|  | NO | 0.0035 | 0.0041 | 1.1714 |
|  | NP | 0.0035 | 0.0058 | 1.6571 |
|  | NR | 0.0035 | 0.0049 | 1.4 |
| *psa*C | NG | 0 | 0.0053 | 0 |
|  | NK | 0 | 0.0053 | 0 |
|  | NO | 0 | 0 | 0 |
|  | NP | 0 | 0.0053 | 0 |
|  | NR | 0 | 0.0052 | 0 |
| *ndh*G | NG | 0 | 0.0023 | 0 |
|  | NK | 0.0104 | 0.0093 | 0.8942 |
|  | NO | 0.0105 | 0.0023 | 0.2190 |
|  | NP | 0.0104 | 0.0117 | 1.1250 |
|  | NR | 0.0103 | 0.0093 | 0.9029 |
| *ndh*I | NG | 0 | 0 | 0 |
|  | NK | 0 | 0 | 0 |
|  | NO | 0 | 0.0051 | 0 |
|  | NP | 0 | 0 | 0 |
|  | NR | 0 | 0 | 0 |
| *ndh*A | NG | 0 | 0.0023 | 0 |
|  | NK | 0.0085 | 0.0046 | 0.5411 |
|  | NO | 0 | 0.0092 | 0 |
|  | NP | 0.0054 | 0 | 0 |
|  | NR | 0.0171 | 0.0046 | 0.2690 |
| *ndh*H | NG | 0 | 0.0021 | 0 |
|  | NK | 0.0043 | 0.0011 | 0.2558 |
|  | NO | 0.0172 | 0.0043 | 0.25 |
|  | NP | 0.0043 | 0.0021 | 0.4883 |
|  | NR | 0.0042 | 0.0021 | 0.5 |
| *rps*15 | NG | 0 | 0 | 0 |
|  | NK | 0 | 0.0144 | 0 |
|  | NO | 0 | 0.0144 | 0 |
|  | NP | 0 | 0.0144 | 0 |
|  | NR | 0 | 0.0142 | 0 |
