## Supplementary Table 5 for "Plastid genomics of *Nicotiana* (Solanaceae): insights into molecular evolution, positive selection and the origin of the maternal genome of Aztec tobacco (*Nicotiana rustica*)"

**Table S5 -** Description of microsatellite loci in the plastid genome of *Nicotiana knightiana*

| **Repeats** | **3** | **4** | **5** | **6** | **7** | **8** | **9** | **10** | **11** | **12** | **13** | **14** | **15** | **16** | **17** | **total** |
| --- | --- | --- | --- | --- | --- | --- | --- | --- | --- | --- | --- | --- | --- | --- | --- | --- |
| A | - | - | - | - | 60 | 23 | 11 | 3 | 2 | 6 | 2 | 1 |  | 1 |  | 109 |
| C | - | - | - | - | 7 | 1 | 1 | 1 |  |  |  |  |  |  |  | 10 |
| G | - | - | - | - | 4 | 1 | 1 |  |  |  |  |  |  |  |  | 6 |
| T | - | - | - | - | 60 | 21 | 22 | 9 | 4 | 1 | 2 |  | 3 | 1 | 1 | 124 |
| AC | - | 1 |  |  |  |  |  |  |  |  |  |  |  |  |  | 1 |
| AG | - | 4 |  |  |  |  |  |  |  |  |  |  |  |  |  | 4 |
| AT | - | 9 | 5 |  |  |  |  |  |  |  |  |  |  |  |  | 14 |
| CT | - | 3 |  |  |  |  |  |  |  |  |  |  |  |  |  | 3 |
| GA | - | 4 |  |  |  |  |  |  |  |  |  |  |  |  |  | 4 |
| TA | - | 9 | 2 |  |  |  |  |  |  |  |  |  |  |  |  | 11 |
| TC | - | 4 |  |  |  |  |  |  |  |  |  |  |  |  |  | 4 |
| AAC | 2 |  |  |  |  |  |  |  |  |  |  |  |  |  |  | 2 |
| AAG | 4 |  |  |  |  |  |  |  |  |  |  |  |  |  |  | 4 |
| AAT | 3 |  |  |  |  |  |  |  |  |  |  |  |  |  |  | 3 |
| AGA | 3 |  |  |  |  |  |  |  |  |  |  |  |  |  |  | 3 |
| ATA | 1 |  |  |  |  |  |  |  |  |  |  |  |  |  |  | 1 |
| ATC | 1 |  |  |  |  |  |  |  |  |  |  |  |  |  |  | 1 |
| ATG | 1 |  |  |  |  |  |  |  |  |  |  |  |  |  |  | 1 |
| ATT | 2 |  |  |  |  |  |  |  |  |  |  |  |  |  |  | 2 |
| CAG | 2 |  |  |  |  |  |  |  |  |  |  |  |  |  |  | 2 |
| CCT | 1 |  |  |  |  |  |  |  |  |  |  |  |  |  |  | 1 |
| CTC | 1 |  |  |  |  |  |  |  |  |  |  |  |  |  |  | 1 |
| CTG | 1 |  |  |  |  |  |  |  |  |  |  |  |  |  |  | 1 |
| CTT | 3 |  |  |  |  |  |  |  |  |  |  |  |  |  |  | 3 |
| GAA | 3 |  |  |  |  |  |  |  |  |  |  |  |  |  |  | 3 |
| GAG | 1 |  |  |  |  |  |  |  |  |  |  |  |  |  |  | 1 |
| GAT | 1 |  |  |  |  |  |  |  |  |  |  |  |  |  |  | 1 |
| GCT | 1 |  |  |  |  |  |  |  |  |  |  |  |  |  |  | 1 |
| GGA | 1 |  |  |  |  |  |  |  |  |  |  |  |  |  |  | 1 |
| GTA | 1 |  |  |  |  |  |  |  |  |  |  |  |  |  |  | 1 |
| GTG | 1 |  |  |  |  |  |  |  |  |  |  |  |  |  |  | 1 |
| GTT | 1 |  |  |  |  |  |  |  |  |  |  |  |  |  |  | 1 |
| TAA | 4 |  |  |  |  |  |  |  |  |  |  |  |  |  |  | 4 |
| TAG | 1 |  |  |  |  |  |  |  |  |  |  |  |  |  |  | 1 |
| TAT | 4 |  |  |  |  |  |  |  |  |  |  |  |  |  |  | 4 |
| TCA | 1 |  |  |  |  |  |  |  |  |  |  |  |  |  |  | 1 |
| TCG | 1 |  |  |  |  |  |  |  |  |  |  |  |  |  |  | 1 |
| TCT | 4 |  |  |  |  |  |  |  |  |  |  |  |  |  |  | 4 |
| TGA | 2 |  |  |  |  |  |  |  |  |  |  |  |  |  |  | 2 |
| TGC | 2 |  |  |  |  |  |  |  |  |  |  |  |  |  |  | 2 |
| TGG | 1 |  |  |  |  |  |  |  |  |  |  |  |  |  |  | 1 |
| TTA | 2 | 2 |  |  |  |  |  |  |  |  |  |  |  |  |  | 4 |
| TTC | 5 | 1 |  |  |  |  |  |  |  |  |  |  |  |  |  | 6 |
| TTG | 4 |  |  |  |  |  |  |  |  |  |  |  |  |  |  | 4 |
| AAAC | 1 |  |  |  |  |  |  |  |  |  |  |  |  |  |  | 1 |
| AAAT | 1 |  |  |  |  |  |  |  |  |  |  |  |  |  |  | 1 |
| CAAA | 1 |  |  |  |  |  |  |  |  |  |  |  |  |  |  | 1 |
| CTAT | 1 |  |  |  |  |  |  |  |  |  |  |  |  |  |  | 1 |
| TAAA | 1 |  |  |  |  |  |  |  |  |  |  |  |  |  |  | 1 |
| TATT | 1 |  |  |  |  |  |  |  |  |  |  |  |  |  |  | 1 |
| TTAT | 1 |  |  |  |  |  |  |  |  |  |  |  |  |  |  | 1 |
| TTTA | 1 |  |  |  |  |  |  |  |  |  |  |  |  |  |  | 1 |
| TTTG | 1 |  |  |  |  |  |  |  |  |  |  |  |  |  |  | 1 |
|  |  |  |  |  |  |  |  |  |  |  |  |  |  |  |  | **368** |

**Table S5a -** Description of microsatellite loci in the plastid genome of *Nicotiana rustica*

| **Repeats** | **3** | **4** | **5** | **6** | **7** | **8** | **9** | **10** | **11** | **12** | **13** | **14** | **15** | **16** | **17** | **total** |
| --- | --- | --- | --- | --- | --- | --- | --- | --- | --- | --- | --- | --- | --- | --- | --- | --- |
| A | - | - | - | - | 61 | 23 | 10 | 2 | 3 | 5 | 1 | 3 |  |  |  | 108 |
| C | - | - | - | - | 7 | 1 | 1 | 1 |  |  |  |  |  |  |  | 10 |
| G | - | - | - | - | 4 | 1 | 1 |  |  |  |  |  |  |  |  | 6 |
| T | - | - | - | - | 60 | 21 | 21 | 11 | 4 | 2 | 1 | 1 | 2 |  | 1 | 124 |
| AC | - | 1 |  |  |  |  |  |  |  |  |  |  |  |  |  | 1 |
| AG | - | 4 |  |  |  |  |  |  |  |  |  |  |  |  |  | 4 |
| AT | - | 9 | 5 |  |  |  |  |  |  |  |  |  |  |  |  | 14 |
| CT | - | 3 |  |  |  |  |  |  |  |  |  |  |  |  |  | 3 |
| GA | - | 4 |  |  |  |  |  |  |  |  |  |  |  |  |  | 4 |
| TA | - | 9 | 2 |  |  |  |  |  |  |  |  |  |  |  |  | 11 |
| TC | - | 4 |  |  |  |  |  |  |  |  |  |  |  |  |  | 4 |
| AAC | 2 |  |  |  |  |  |  |  |  |  |  |  |  |  |  | 2 |
| AAG | 4 |  |  |  |  |  |  |  |  |  |  |  |  |  |  | 4 |
| AAT | 3 |  |  |  |  |  |  |  |  |  |  |  |  |  |  | 3 |
| AGA | 3 |  |  |  |  |  |  |  |  |  |  |  |  |  |  | 3 |
| ATA | 1 |  |  |  |  |  |  |  |  |  |  |  |  |  |  | 1 |
| ATC | 1 |  |  |  |  |  |  |  |  |  |  |  |  |  |  | 1 |
| ATG | 1 |  |  |  |  |  |  |  |  |  |  |  |  |  |  | 1 |
| ATT | 2 |  |  |  |  |  |  |  |  |  |  |  |  |  |  | 2 |
| CAG | 2 |  |  |  |  |  |  |  |  |  |  |  |  |  |  | 2 |
| CCT | 1 |  |  |  |  |  |  |  |  |  |  |  |  |  |  | 1 |
| CTC | 1 |  |  |  |  |  |  |  |  |  |  |  |  |  |  | 1 |
| CTG | 1 |  |  |  |  |  |  |  |  |  |  |  |  |  |  | 1 |
| CTT | 3 |  |  |  |  |  |  |  |  |  |  |  |  |  |  | 3 |
| GAA | 3 |  |  |  |  |  |  |  |  |  |  |  |  |  |  | 3 |
| GAG | 1 |  |  |  |  |  |  |  |  |  |  |  |  |  |  | 1 |
| GAT | 1 |  |  |  |  |  |  |  |  |  |  |  |  |  |  | 1 |
| GCT | 1 |  |  |  |  |  |  |  |  |  |  |  |  |  |  | 1 |
| GGA | 1 |  |  |  |  |  |  |  |  |  |  |  |  |  |  | 1 |
| GTA | 1 |  |  |  |  |  |  |  |  |  |  |  |  |  |  | 1 |
| GTG | 1 |  |  |  |  |  |  |  |  |  |  |  |  |  |  | 1 |
| GTT | 1 |  |  |  |  |  |  |  |  |  |  |  |  |  |  | 1 |
| TAA | 4 |  |  |  |  |  |  |  |  |  |  |  |  |  |  | 4 |
| TAG | 1 |  |  |  |  |  |  |  |  |  |  |  |  |  |  | 1 |
| TAT | 4 |  |  |  |  |  |  |  |  |  |  |  |  |  |  | 4 |
| TCA | 1 |  |  |  |  |  |  |  |  |  |  |  |  |  |  | 1 |
| TCG | 1 |  |  |  |  |  |  |  |  |  |  |  |  |  |  | 1 |
| TCT | 4 |  |  |  |  |  |  |  |  |  |  |  |  |  |  | 4 |
| TGA | 2 |  |  |  |  |  |  |  |  |  |  |  |  |  |  | 2 |
| TGC | 2 |  |  |  |  |  |  |  |  |  |  |  |  |  |  | 2 |
| TGG | 1 |  |  |  |  |  |  |  |  |  |  |  |  |  |  | 1 |
| TTA | 2 | 2 |  |  |  |  |  |  |  |  |  |  |  |  |  | 4 |
| TTC | 5 | 1 |  |  |  |  |  |  |  |  |  |  |  |  |  | 6 |
| TTG | 4 |  |  |  |  |  |  |  |  |  |  |  |  |  |  | 4 |
| AAAC | 1 |  |  |  |  |  |  |  |  |  |  |  |  |  |  | 1 |
| AAAT | 1 |  |  |  |  |  |  |  |  |  |  |  |  |  |  | 1 |
| CAAA | 1 |  |  |  |  |  |  |  |  |  |  |  |  |  |  | 1 |
| CTAT | 1 |  |  |  |  |  |  |  |  |  |  |  |  |  |  | 1 |
| TAAA | 1 |  |  |  |  |  |  |  |  |  |  |  |  |  |  | 1 |
| TATT | 1 |  |  |  |  |  |  |  |  |  |  |  |  |  |  | 1 |
| TTAT | 1 |  |  |  |  |  |  |  |  |  |  |  |  |  |  | 1 |
| TTTA | 1 |  |  |  |  |  |  |  |  |  |  |  |  |  |  | 1 |
| TTTG | 1 |  |  |  |  |  |  |  |  |  |  |  |  |  |  | 1 |
|  |  |  |  |  |  |  |  |  |  |  |  |  |  |  |  | 367 |

**Table S5b -** Description of microsatellite loci in the plastid genome of *Nicotiana paniculata*

| **Repeats** | **3** | **4** | **5** | **6** | **7** | **8** | **9** | **10** | **11** | **12** | **13** | **14** | **15** | **16** | **17** | **18** | **total** |
| --- | --- | --- | --- | --- | --- | --- | --- | --- | --- | --- | --- | --- | --- | --- | --- | --- | --- |
| A | - | - | - | - | 61 | 23 | 10 | 3 | 4 | 1 | 2 |  | 2 | 1 |  |  | 107 |
| C | - | - | - | - | 7 | 1 |  |  |  | 1 |  | 1 |  |  |  |  | 10 |
| G | - | - | - | - | 4 | 1 | 1 |  |  |  |  |  |  |  |  |  | 6 |
| T | - | - | - | - | 60 | 21 | 25 | 9 | 3 | 3 |  |  | 3 | 1 |  | 1 | 126 |
| AC | - | 1 |  |  |  |  |  |  |  |  |  |  |  |  |  |  | 1 |
| AG | - | 5 |  |  |  |  |  |  |  |  |  |  |  |  |  |  | 5 |
| AT | - | 9 | 5 |  |  |  |  |  |  |  |  |  |  |  |  |  | 14 |
| CT | - | 3 |  |  |  |  |  |  |  |  |  |  |  |  |  |  | 3 |
| GA | - | 4 |  |  |  |  |  |  |  |  |  |  |  |  |  |  | 4 |
| TA | - | 9 | 2 |  |  |  |  |  |  |  |  |  |  |  |  |  | 11 |
| TC | - | 4 |  |  |  |  |  |  |  |  |  |  |  |  |  |  | 4 |
| AAC | 2 |  |  |  |  |  |  |  |  |  |  |  |  |  |  |  | 2 |
| AAG | 4 |  |  |  |  |  |  |  |  |  |  |  |  |  |  |  | 4 |
| AAT | 3 |  |  |  |  |  |  |  |  |  |  |  |  |  |  |  | 3 |
| AGA | 3 |  |  |  |  |  |  |  |  |  |  |  |  |  |  |  | 3 |
| ATA | 1 |  |  |  |  |  |  |  |  |  |  |  |  |  |  |  | 1 |
| ATC | 1 |  |  |  |  |  |  |  |  |  |  |  |  |  |  |  | 1 |
| ATG | 1 |  |  |  |  |  |  |  |  |  |  |  |  |  |  |  | 1 |
| ATT | 2 |  |  |  |  |  |  |  |  |  |  |  |  |  |  |  | 2 |
| CAG | 2 |  |  |  |  |  |  |  |  |  |  |  |  |  |  |  | 2 |
| CCT | 1 |  |  |  |  |  |  |  |  |  |  |  |  |  |  |  | 1 |
| CTA | 1 |  |  |  |  |  |  |  |  |  |  |  |  |  |  |  | 1 |
| CTC | 1 |  |  |  |  |  |  |  |  |  |  |  |  |  |  |  | 1 |
| CTG | 1 |  |  |  |  |  |  |  |  |  |  |  |  |  |  |  | 1 |
| CTT | 3 |  |  |  |  |  |  |  |  |  |  |  |  |  |  |  | 3 |
| GAA | 3 |  |  |  |  |  |  |  |  |  |  |  |  |  |  |  | 3 |
| GAG | 1 |  |  |  |  |  |  |  |  |  |  |  |  |  |  |  | 1 |
| GAT | 1 |  |  |  |  |  |  |  |  |  |  |  |  |  |  |  | 1 |
| GCT | 1 |  |  |  |  |  |  |  |  |  |  |  |  |  |  |  | 1 |
| GGA | 1 |  |  |  |  |  |  |  |  |  |  |  |  |  |  |  | 1 |
| GTA | 1 |  |  |  |  |  |  |  |  |  |  |  |  |  |  |  | 1 |
| GTG | 1 |  |  |  |  |  |  |  |  |  |  |  |  |  |  |  | 1 |
| GTT | 1 |  |  |  |  |  |  |  |  |  |  |  |  |  |  |  | 1 |
| TAA | 4 |  |  |  |  |  |  |  |  |  |  |  |  |  |  |  | 4 |
| TAG | 1 |  |  |  |  |  |  |  |  |  |  |  |  |  |  |  | 1 |
| TAT | 3 |  |  |  |  |  |  |  |  |  |  |  |  |  |  |  | 3 |
| TCA | 1 |  |  |  |  |  |  |  |  |  |  |  |  |  |  |  | 1 |
| TCG | 1 |  |  |  |  |  |  |  |  |  |  |  |  |  |  |  | 1 |
| TCT | 4 |  |  |  |  |  |  |  |  |  |  |  |  |  |  |  | 4 |
| TGA | 2 |  |  |  |  |  |  |  |  |  |  |  |  |  |  |  | 2 |
| TGC | 2 |  |  |  |  |  |  |  |  |  |  |  |  |  |  |  | 2 |
| TGG | 1 |  |  |  |  |  |  |  |  |  |  |  |  |  |  |  | 1 |
| TTA | 2 | 2 |  |  |  |  |  |  |  |  |  |  |  |  |  |  | 4 |
| TTC | 6 | 1 |  |  |  |  |  |  |  |  |  |  |  |  |  |  | 7 |
| TTG | 4 |  |  |  |  |  |  |  |  |  |  |  |  |  |  |  | 4 |
| AAAC | 1 |  |  |  |  |  |  |  |  |  |  |  |  |  |  |  | 1 |
| AAAT | 1 |  |  |  |  |  |  |  |  |  |  |  |  |  |  |  | 1 |
| CAAA | 1 |  |  |  |  |  |  |  |  |  |  |  |  |  |  |  | 1 |
| CTAT | 1 |  |  |  |  |  |  |  |  |  |  |  |  |  |  |  | 1 |
| TAAA | 1 |  |  |  |  |  |  |  |  |  |  |  |  |  |  |  | 1 |
| TATT | 1 |  |  |  |  |  |  |  |  |  |  |  |  |  |  |  | 1 |
| TTAT | 1 |  |  |  |  |  |  |  |  |  |  |  |  |  |  |  | 1 |
| TTTA | 1 |  |  |  |  |  |  |  |  |  |  |  |  |  |  |  | 1 |
| TTTG | 1 |  |  |  |  |  |  |  |  |  |  |  |  |  |  |  | 1 |
|  |  |  |  |  |  |  |  |  |  |  |  |  |  |  |  |  | 370 |

**Table S5c -** Description of microsatellite loci in the plastid genome of *Nicotiana obtusifolia*

| **Repeats** | **3** | **4** | **5** | **6** | **7** | **8** | **9** | **10** | **11** | **12** | **13** | **14** | **15** | **16** | **17** | **18** | **total** |
| --- | --- | --- | --- | --- | --- | --- | --- | --- | --- | --- | --- | --- | --- | --- | --- | --- | --- |
| A | - | - | - | - | 66 | 24 | 11 | 7 | 5 | 2 |  | 1 |  | 1 |  |  | 117 |
| C | - | - | - | - | 9 | 3 |  |  |  |  |  |  |  |  |  |  | 12 |
| G | - | - | - | - | 5 | 1 | 1 |  |  |  |  |  |  |  |  |  | 7 |
| T | - | - | - | - | 63 | 23 | 13 | 11 | 11 | 2 | 3 | 1 |  |  |  | 1 | 128 |
| AC | - | 1 |  |  |  |  |  |  |  |  |  |  |  |  |  |  | 1 |
| AG | - | 5 |  |  |  |  |  |  |  |  |  |  |  |  |  |  | 5 |
| AT | - | 8 | 5 |  | 1 |  |  |  |  |  |  |  |  |  |  |  | 14 |
| CT | - | 3 |  |  |  |  |  |  |  |  |  |  |  |  |  |  | 3 |
| GA | - | 4 |  |  |  |  |  |  |  |  |  |  |  |  |  |  | 4 |
| TA | - | 10 | 2 |  |  |  |  |  |  |  |  |  |  |  |  |  | 12 |
| TC | - | 4 |  |  |  |  |  |  |  |  |  |  |  |  |  |  | 4 |
| AAC | 2 |  |  |  |  |  |  |  |  |  |  |  |  |  |  |  | 2 |
| AAG | 3 | 1 |  |  |  |  |  |  |  |  |  |  |  |  |  |  | 4 |
| AAT | 4 |  |  |  |  |  |  |  |  |  |  |  |  |  |  |  | 4 |
| AGA | 3 |  |  |  |  |  |  |  |  |  |  |  |  |  |  |  | 3 |
| ATA | 1 |  |  |  |  |  |  |  |  |  |  |  |  |  |  |  | 1 |
| ATC | 1 |  |  |  |  |  |  |  |  |  |  |  |  |  |  |  | 1 |
| ATG | 1 |  |  |  |  |  |  |  |  |  |  |  |  |  |  |  | 1 |
| ATT | 2 | 1 |  |  |  |  |  |  |  |  |  |  |  |  |  |  | 3 |
| CAG | 2 |  |  |  |  |  |  |  |  |  |  |  |  |  |  |  | 2 |
| CCT | 1 |  |  |  |  |  |  |  |  |  |  |  |  |  |  |  | 1 |
| CTC | 1 |  |  |  |  |  |  |  |  |  |  |  |  |  |  |  | 1 |
| CTG | 1 |  |  |  |  |  |  |  |  |  |  |  |  |  |  |  | 1 |
| CTT | 3 |  |  |  |  |  |  |  |  |  |  |  |  |  |  |  | 3 |
| GAA | 3 |  |  |  |  |  |  |  |  |  |  |  |  |  |  |  | 3 |
| GAG | 1 |  |  |  |  |  |  |  |  |  |  |  |  |  |  |  | 1 |
| GAT | 1 |  |  |  |  |  |  |  |  |  |  |  |  |  |  |  | 1 |
| GCT | 1 |  |  |  |  |  |  |  |  |  |  |  |  |  |  |  | 1 |
| GGA | 1 |  |  |  |  |  |  |  |  |  |  |  |  |  |  |  | 1 |
| GTA | 1 |  |  |  |  |  |  |  |  |  |  |  |  |  |  |  | 1 |
| GTG | 1 |  |  |  |  |  |  |  |  |  |  |  |  |  |  |  | 1 |
| GTT | 1 |  |  |  |  |  |  |  |  |  |  |  |  |  |  |  | 1 |
| TAA | 4 |  |  |  |  |  |  |  |  |  |  |  |  |  |  |  | 4 |
| TAG | 1 |  |  |  |  |  |  |  |  |  |  |  |  |  |  |  | 1 |
| TAT | 3 |  |  |  |  |  |  |  |  |  |  |  |  |  |  |  | 3 |
| TCA | 1 |  |  |  |  |  |  |  |  |  |  |  |  |  |  |  | 1 |
| TCG | 1 |  |  |  |  |  |  |  |  |  |  |  |  |  |  |  | 1 |
| TCT | 4 |  |  |  |  |  |  |  |  |  |  |  |  |  |  |  | 4 |
| TGA | 2 |  |  |  |  |  |  |  |  |  |  |  |  |  |  |  | 2 |
| TGC | 2 |  |  |  |  |  |  |  |  |  |  |  |  |  |  |  | 2 |
| TGG | 1 |  |  |  |  |  |  |  |  |  |  |  |  |  |  |  | 1 |
| TTA | 2 | 1 |  |  |  |  |  |  |  |  |  |  |  |  |  |  | 3 |
| TTC | 5 | 1 |  |  |  |  |  |  |  |  |  |  |  |  |  |  | 6 |
| TTG | 4 |  |  |  |  |  |  |  |  |  |  |  |  |  |  |  | 4 |
| AAAC | 1 |  |  |  |  |  |  |  |  |  |  |  |  |  |  |  | 1 |
| AAAT | 1 |  |  |  |  |  |  |  |  |  |  |  |  |  |  |  | 1 |
| CAAA | 1 |  |  |  |  |  |  |  |  |  |  |  |  |  |  |  | 1 |
| CTAT | 1 |  |  |  |  |  |  |  |  |  |  |  |  |  |  |  | 1 |
| TAAA | 1 |  |  |  |  |  |  |  |  |  |  |  |  |  |  |  | 1 |
| TTTA | 1 |  |  |  |  |  |  |  |  |  |  |  |  |  |  |  | 1 |
| TTTG | 1 |  |  |  |  |  |  |  |  |  |  |  |  |  |  |  | 1 |
| TTTAA | 1 |  |  |  |  |  |  |  |  |  |  |  |  |  |  |  | 1 |
|  |  |  |  |  |  |  |  |  |  |  |  |  |  |  |  |  | 384 |

**Table S5d -** Description of microsatellite loci in the plastid genome of *Nicotiana glauca*

| **Repeats** | **3** | **4** | **5** | **6** | **7** | **8** | **9** | **10** | **11** | **12** | **13** | **14** | **15** | **16** | **17** | **18** | **19** | **20** | **21** | **total** |
| --- | --- | --- | --- | --- | --- | --- | --- | --- | --- | --- | --- | --- | --- | --- | --- | --- | --- | --- | --- | --- |
| A | - | - | - | - | 65 | 23 | 11 | 7 | 1 | 1 |  | 1 | 1 |  | 1 |  |  |  |  | 111 |
| C | - | - | - | - | 7 | 1 |  |  | 1 |  |  |  |  |  |  |  |  |  |  | 9 |
| G | - | - | - | - | 5 | 2 |  |  |  |  |  |  |  |  |  |  |  |  |  | 7 |
| T | - | - | - | - | 60 | 24 | 17 | 9 | 6 | 3 |  | 3 | 1 | 1 |  |  |  |  | 1 | 125 |
| AC | - | 1 |  |  |  |  |  |  |  |  |  |  |  |  |  |  |  |  |  | 1 |
| AG | - | 5 |  |  |  |  |  |  |  |  |  |  |  |  |  |  |  |  |  | 5 |
| AT | - | 11 | 5 |  |  |  |  |  |  |  |  |  |  |  |  |  |  |  |  | 16 |
| CT | - | 2 |  |  |  |  |  |  |  |  |  |  |  |  |  |  |  |  |  | 2 |
| GA | - | 4 |  |  |  |  |  |  |  |  |  |  |  |  |  |  |  |  |  | 4 |
| TA | - | 9 | 2 |  |  |  |  |  |  |  |  |  |  |  |  |  |  |  |  | 11 |
| TC | - | 4 |  |  |  |  |  |  |  |  |  |  |  |  |  |  |  |  |  | 4 |
| AAC | 2 |  |  |  |  |  |  |  |  |  |  |  |  |  |  |  |  |  |  | 2 |
| AAG | 3 | 1 |  |  |  |  |  |  |  |  |  |  |  |  |  |  |  |  |  | 4 |
| AAT | 3 |  |  |  |  |  |  |  |  |  |  |  |  |  |  |  |  |  |  | 3 |
| AGA | 3 |  |  |  |  |  |  |  |  |  |  |  |  |  |  |  |  |  |  | 3 |
| ATA | 1 | 1 |  |  |  |  |  |  |  |  |  |  |  |  |  |  |  |  |  | 2 |
| ATC | 1 |  |  |  |  |  |  |  |  |  |  |  |  |  |  |  |  |  |  | 1 |
| ATG | 1 |  |  |  |  |  |  |  |  |  |  |  |  |  |  |  |  |  |  | 1 |
| ATT | 2 |  |  |  |  |  |  |  |  |  |  |  |  |  |  |  |  |  |  | 2 |
| CAG | 2 |  |  |  |  |  |  |  |  |  |  |  |  |  |  |  |  |  |  | 2 |
| CCT | 1 |  |  |  |  |  |  |  |  |  |  |  |  |  |  |  |  |  |  | 1 |
| CTC | 1 |  |  |  |  |  |  |  |  |  |  |  |  |  |  |  |  |  |  | 1 |
| CTG | 1 |  |  |  |  |  |  |  |  |  |  |  |  |  |  |  |  |  |  | 1 |
| CTT | 3 |  |  |  |  |  |  |  |  |  |  |  |  |  |  |  |  |  |  | 3 |
| GAA | 3 |  |  |  |  |  |  |  |  |  |  |  |  |  |  |  |  |  |  | 3 |
| GAG | 1 |  |  |  |  |  |  |  |  |  |  |  |  |  |  |  |  |  |  | 1 |
| GAT | 1 |  |  |  |  |  |  |  |  |  |  |  |  |  |  |  |  |  |  | 1 |
| GCT | 1 |  |  |  |  |  |  |  |  |  |  |  |  |  |  |  |  |  |  | 1 |
| GGA | 1 |  |  |  |  |  |  |  |  |  |  |  |  |  |  |  |  |  |  | 1 |
| GTA | 1 |  |  |  |  |  |  |  |  |  |  |  |  |  |  |  |  |  |  | 1 |
| GTG | 1 |  |  |  |  |  |  |  |  |  |  |  |  |  |  |  |  |  |  | 1 |
| GTT | 1 |  |  |  |  |  |  |  |  |  |  |  |  |  |  |  |  |  |  | 1 |
| TAA | 4 |  |  |  |  |  |  |  |  |  |  |  |  |  |  |  |  |  |  | 4 |
| TAG | 1 |  |  |  |  |  |  |  |  |  |  |  |  |  |  |  |  |  |  | 1 |
| TAT | 3 |  |  |  |  |  |  |  |  |  |  |  |  |  |  |  |  |  |  | 3 |
| TCA | 1 |  |  |  |  |  |  |  |  |  |  |  |  |  |  |  |  |  |  | 1 |
| TCG | 1 |  |  |  |  |  |  |  |  |  |  |  |  |  |  |  |  |  |  | 1 |
| TCT | 4 |  |  |  |  |  |  |  |  |  |  |  |  |  |  |  |  |  |  | 4 |
| TGA | 2 |  |  |  |  |  |  |  |  |  |  |  |  |  |  |  |  |  |  | 2 |
| TGC | 2 |  |  |  |  |  |  |  |  |  |  |  |  |  |  |  |  |  |  | 2 |
| TGG | 1 |  |  |  |  |  |  |  |  |  |  |  |  |  |  |  |  |  |  | 1 |
| TTA | 2 | 2 |  |  |  |  |  |  |  |  |  |  |  |  |  |  |  |  |  | 4 |
| TTC | 6 | 1 |  |  |  |  |  |  |  |  |  |  |  |  |  |  |  |  |  | 7 |
| TTG | 4 |  |  |  |  |  |  |  |  |  |  |  |  |  |  |  |  |  |  | 4 |
| AAAC | 1 |  |  |  |  |  |  |  |  |  |  |  |  |  |  |  |  |  |  | 1 |
| CAAA | 1 |  |  |  |  |  |  |  |  |  |  |  |  |  |  |  |  |  |  | 1 |
| CTAT | 1 |  |  |  |  |  |  |  |  |  |  |  |  |  |  |  |  |  |  | 1 |
| TAAA | 1 |  |  |  |  |  |  |  |  |  |  |  |  |  |  |  |  |  |  | 1 |
| TATT | 1 |  |  |  |  |  |  |  |  |  |  |  |  |  |  |  |  |  |  | 1 |
| TTTA | 1 |  |  |  |  |  |  |  |  |  |  |  |  |  |  |  |  |  |  | 1 |
| TTTG | 1 |  |  |  |  |  |  |  |  |  |  |  |  |  |  |  |  |  |  | 371 |
